# PmeR, a TetR-like transcriptional regulator, is involved in both auxin signaling and virulence in the plant pathogen *Pseudomonas syringae* strain *Pto*DC3000

**DOI:** 10.1101/2025.03.31.646273

**Authors:** Chia-Yun Lee, Maya Irvine, Barbara Kunkel

## Abstract

Plant pathogenic bacteria, such as *Pseudomonas syringae* strain *Pto*DC3000, respond to host signals through complex signaling networks that regulate bacterial growth and virulence. The plant hormone indole-3-acetic acid (IAA), also known as auxin, promotes bacterial pathogenesis via multiple mechanisms, including through reprogramming bacterial transcription. However, the mechanisms that *Pto*DC3000 uses to sense and respond to auxin are not well understood. Here, we identify *pmeR*, which encodes a TetR-like family transcriptional repressor, as an important regulator of IAA-responsive gene expression in *Pto*DC3000. Using qRT-PCR and transcriptional reporter assays, we show that *pmeR* is induced by IAA and regulates a set of auxin-responsive genes, including itself as well as several genes known or proposed to be involved in virulence. Plant infection assays further show that the disruption of *pmeR* results in reduced bacterial growth in *Arabidopsis thaliana*. Notably, while PmeR de-represses transcription of *pmeR* upon IAA treatment, it does not appear to directly bind IAA. Rather, our biochemical results indicate that the auxin conjugate IAA-Lysine may serve as a ligand for PmeR. Our findings reveal a complex signaling network through which IAA modulates bacterial gene expression and emphasizes the role of PmeR in acclimating *Pto*DC3000 for growth in plant tissue.

**Author Summary:** Plant pathogenic bacteria, such as *Pseudomonas syringae* strain *Pto*DC3000, respond to host signals through complex signaling networks that regulate bacterial growth and virulence. One key signal involved in these interactions is the plant hormone indole-3-acetic acid (IAA), which has been shown to promote bacterial pathogenicity in *Arabidopsis thaliana* and tomato. However, the mechanisms *Pto*DC3000 uses to sense and respond to IAA remain poorly understood. In this study, we explored the role of the TetR-like transcriptional regulator PmeR, encoded by the *pmeR* gene, in regulating bacterial responses to IAA. We found that *pmeR* is induced by IAA and regulates the expression of several additional auxin-responsive genes. Furthermore, we showed that *pmeR* is required for full virulence of *Pto*DC3000 in *A. thaliana*. These results suggest that PmeR is involved in regulating IAA-induced gene expression and that the ability to respond to auxin contributes to virulence of *Pto*DC3000. This work sheds new light on the molecular mechanisms through which IAA regulates bacterial pathogenesis, providing important insights into plant-microbe interactions and the role of auxin in regulating bacterial behavior. Our findings offer potential directions for developing strategies to mitigate bacterial diseases in crops by targeting auxin-responsive regulatory pathways.

## Introduction

Microorganisms rely on complex signaling mechanisms to enable them to adjust their physiology and metabolism to changing environmental conditions. Included among known signaling molecules (1–4) is the plant hormone auxin (indole-3-acetic acid, IAA), which plays a critical role in plant-pathogen interactions by both promoting plant host susceptibility and modulating bacterial pathogenesis (5–7).

*Pseudomonas syringae* strain *Pto*DC3000 (*Pto*DC3000) is a hemibiotrophic phytopathogenic bacterium that infects both tomato (*Solanum lycopersicon*) and *Arabidopsis thaliana*. Previously, we demonstrated that in *Pto*DC3000, IAA promotes its pathogenesis in *A. thaliana* by two different mechanisms: (i) suppressing salicylic acid-mediated basal host defenses and (ii) regulating bacterial virulence gene expression (8, 9). We also observed that several of the IAA-induced genes are predicted to encode transcriptional regulators (10), leading us to hypothesize that one or more of these regulators might be involved in regulating the IAA response in *Pto*DC3000. In this study, we investigated the role of one of these IAA-induced regulatory genes, *pmeR*, which encodes a TetR-like family transcriptional regulator, in regulating IAA responsiveness.

TetR-like family transcriptional regulators (TFRs) allow bacteria to respond to changing environmental conditions by sensing small molecules inside the cell (11). TFRs typically function as transcriptional repressors that bind to palindromic DNA sequences in the regulatory regions (e.g., the operator) of their own structural gene, thus repressing their own expression. Often these operator sequences also regulate expression of an adjacent operon that encodes an efflux pump. The N-terminal domains of TFRs, which feature a conserved helix-turn-helix (HTH) motif, are responsible for specific binding to the operator DNA. Meanwhile, the C-terminal domains are required for dimerization and ligand binding. The ligands are often small molecules that are also substrates of the efflux pumps encoded by the adjacent operons. In the absence of their ligand, TFRs maintain a default conformation that allows them to bind to the operator, preventing RNA polymerase from binding to the promoter and thus repressing transcription of both their own genes and adjacent operons. Upon binding a specific ligand, the TFR undergoes a conformational change, causing the ligand-protein complex to be released from the operator and de-repressing gene expression (11).

In *Pto*DC3000, PmeR represses both its own gene (*pmeR, PSPTO_4302*) and the divergently oriented *mexAB-oprM* operon (*PSPO_4303*-*4305*), which encodes a resistance-nodulation-division (RND) multidrug efflux system (12). Plant-derived flavonoids, such as phloretin, have been shown to cause the dissociation of PmeR from the operator within the intergenic regulatory region, resulting in de-repression of both *pmeR* and the *mexAB-oprM* operon and subsequently conferring increased antimicrobial tolerance to flavonoids and other antibiotics. Here, we demonstrate that *pmeR* is induced by IAA and regulates a set of auxin-responsive genes, including its own gene and several known or proposed to be involved in virulence. Additionally, we show that *pmeR* contributes to *Pto*DC3000 virulence in *Arabidopsis*. Our biochemical assays further demonstrate that rather than IAA, the IAA-amino acid conjugate IAA-Lysine impacts the binding between PmeR and its operator. This reveals that IAA perception in *Pto*DC3000 is mediated via a complex signaling network and contributes to pathogen virulence.

## Results

### *pmeR* is induced by IAA and regulates the sensitivity of *Pto*DC3000 to IAA

Transcription of both *pmeR* and the adjacent *mexAB-oprM* operon, which is directly regulated by *pmeR* (12), was previously shown to be upregulated by IAA in culture (10). To confirm this, we monitored the expression of *pmeR* and *mexA* in *Pto*DC3000 cells grown in Hrp de-repressing medium (HDM) with or without 100 µM IAA, using quantitative reverse transcription polymerase chain reaction (qRT-PCR). As shown in Figs 1A and 1B, we confirmed that IAA significantly upregulates transcript levels of both *pmeR* and *mexA* at 30 minutes post-treatment.

**Fig 1.**
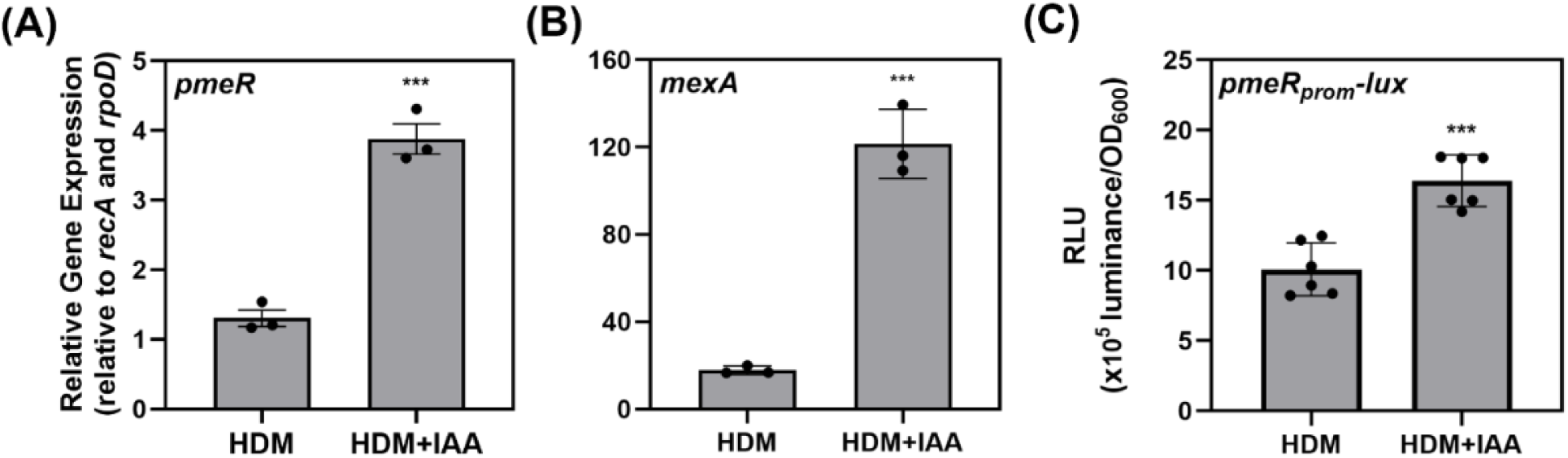
*pmeR* and *mexA* are upregulated by IAA in culture. Expression of (A) *pmeR* (*PSPTO_4302*) and (B) *mexA* (*PSPTO_4303*) in wild-type *Pto*DC3000 (WT) in response to IAA at 30 minutes post-treatment was monitored by qRT-PCR. Data represent mean expression normalized to *rpoD* (*PSPTO_0537*) and *recA* (*PSPTO_4033*), plotted as relative expression ± standard deviation (SD). Data are from a single representative experiment (n = 3). Similar results were obtained in a second independent experiment. (C) Expression of *pmeR_prom_-lux* in WT was analyzed by quantifying luminescence at 90 minutes post-treatment. Data represent the mean of relative luminescence normalized to cell density (OD_600_), plotted as relative luminescence units (RLU) ± SD. Data were compiled from two independent experiments (n = 6). Asterisks indicate significant differences between treatments as determined by Student’s *t*-test (***: *p* < 0.001). HDM: Hrp de-repressing medium; HDM + IAA: HDM supplemented with 100 µM IAA.

PmeR is a transcriptional repressor that de-represses the expression of its own gene and the divergently oriented gene operon in response to intracellular signals (12). Given that both *pmeR* and *mexA* are upregulated by IAA, we investigated whether *pmeR* might regulate its own auxin-responsiveness. We generated a transcriptional fusion to monitor the promoter activity of *pmeR*, and introduced it into wild-type *Pto*DC3000 (WT) and a previously described *Pto*DC3000 *pmeR*::ΩKan mutant (12) (hereafter referred to as “*pmeR”*). The fusion consisted of the bacterial luciferase operon from *Photorhabdus luminescens* driven by the promoter of the *pmeR* gene (*pmeR_prom_-lux*), carried on a stable plasmid (S1 Table) (13–15). Basal expression of *pmeR_prom_-lux* was significantly higher in the *pmeR* mutant than in WT cells (S1 Fig), consistent with previous findings that PmeR acts as a transcriptional repressor of *pmeR* (12). We also used the *pmeR_prom_- lux* reporter to confirm that IAA induces expression of *pmeR_prom_-lux* in WT (Fig 1C), demonstrating that this transcriptional fusion is a reliable reporter of *pmeR* expression.

We next utilized the *pmeR_prom_-lux* fusion to examine if *pmeR* might regulate the IAA- responsiveness of its own expression by monitoring the level of IAA-induction of *pmeR_prom_-lux* expression in WT and *pmeR* mutant at 30-, 60-, and 90-minutes post-treatment. The addition of IAA caused a slight but insignificant increase in *pmeR_prom_-lux* expression in WT cells at 30 minutes post-treatment, and this increase became significant by 60 minutes post-treatment (Fig 2). Notably, at 60 minutes post-treatment, the increased level of expression in WT was similar to the de-repressed level observed in the *pmeR* mutant in the absence of IAA. On the other hand, in the *pmeR* mutant, the addition of IAA did not significantly induce expression of *pmeR_prom_-lux* beyond the de-repressed level at this time point. However, by 90 minutes, the addition of IAA induced expression of *pmeR_prom_-lux* to above de-repressed levels in both WT and *pmeR* mutant cells (Fig 2). These observations indicate that induction of *pmeR* by IAA is dependent upon *pmeR* at 60 minutes but is independent of *pmeR* by 90 minutes, suggesting additional transcriptional regulators contribute to auxin induction at the later time point (90 minutes post-treatment).

**Fig 2.**
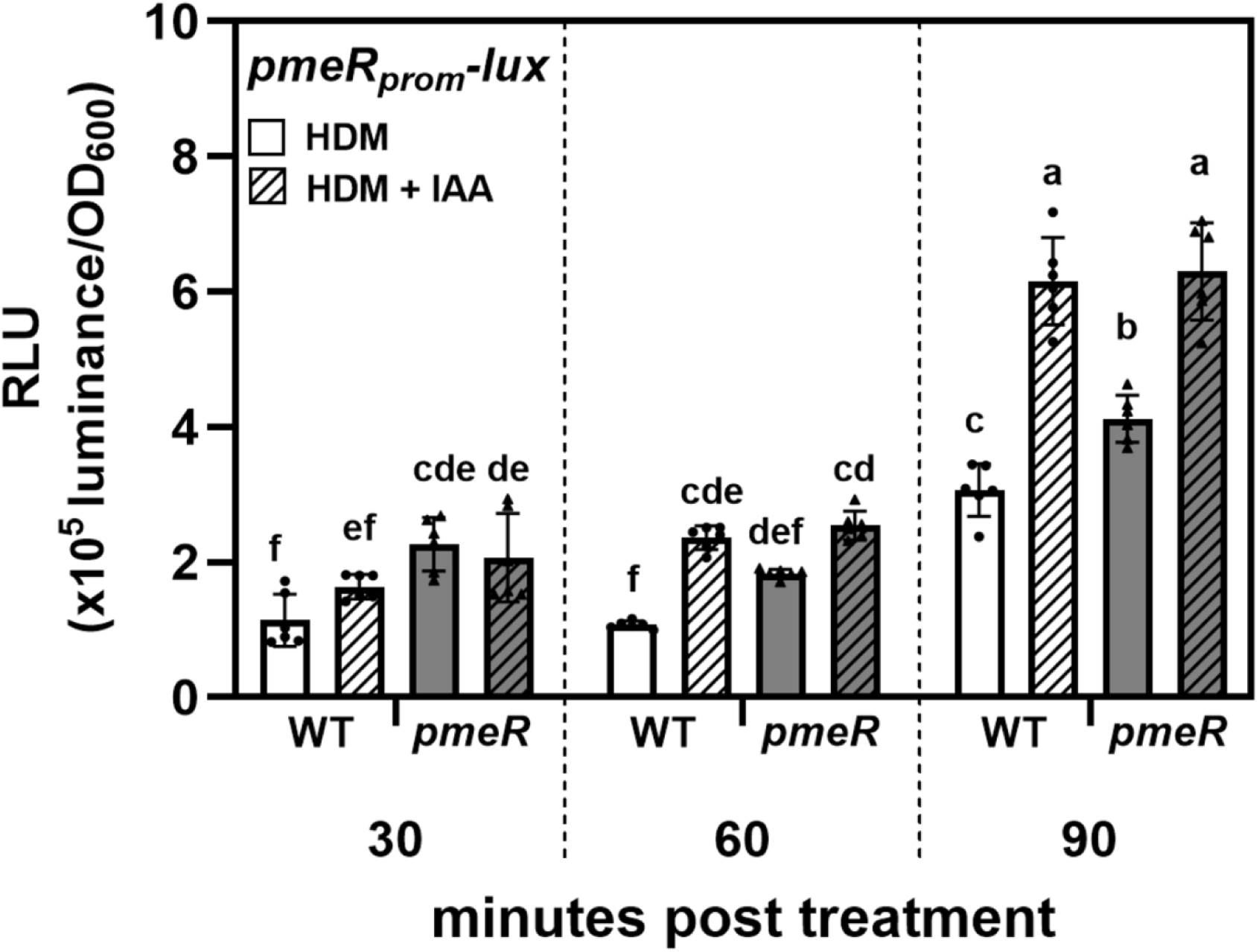
Mutation of *pmeR* reduces *Pto*DC3000 sensitivity to IAA. Expression of the *pmeR_prom_- lux* reporter in wild-type *Pto*DC3000 (WT) and the *pmeR*::ΩKan mutant (*pmeR*) in response to IAA at indicated time points post-treatment with IAA. Data represent mean luminescence normalized to OD_600_ and are plotted as the mean of relative luminescence units (RLU) ± SD. Data are compiled from two independent experiments (n = 6). Letters indicate significant differences between treatments as determined by ANOVA followed by Tukey’s HSD test (*p* < 0.05). HDM: Hrp de-repressing medium; HDM + IAA: HDM supplemented with 100 µM IAA.

### *pmeR* regulates the expression of several additional auxin-responsive genes in *Pto*DC3000

Given that *pmeR* encodes an auxin-upregulated transcription factor, we hypothesized that it may regulate the expression of other auxin-responsive genes in addition to *pmeR*. To test this, we selected a set of genes previously identified as auxin-responsive (either auxin-upregulated or auxin-downregulated) that are also expressed *in planta* (2, 4) and quantified their transcript levels in WT and the *pmeR* mutant in response to IAA in culture. These genes included *PSPTO_1824* and *PSPTO_4297*, both predicted to encode small conserved hypothetical proteins of unknown function, and *PSPTO_3549* (*aefR*) and *PSPTO_3576* (*tvrR*), which encode transcription factors that contribute to virulence (15, 16). *Pto*DC3000 cells were grown to mid-log phase in nutrient yeast glycerol medium (NYG), transferred to HDM supplemented with or without 100 µM IAA, and harvested at 30 minutes post-treatment for qRT-PCR analysis. As expected, the transcript levels of these four auxin-upregulated genes, *PSPTO_1824*, *PSPTO_4297*, *PSPTO_3549* (*aefR*), and *PSPTO_3576* (*tvrR*), were strongly induced by IAA in WT cells compared to the low expression levels observed in NYG and HDM media (Figs 3A-3D). The expression level of three of these genes, *PSPTO_1824*, *PSPTO_4297*, and *PSPTO_3549* (*aefR*), in response to auxin was significantly reduced in the *pmeR* mutant compared to WT, indicating that up-regulation of these genes by IAA is partially dependent on *pmeR*. In contrast, the level of IAA induction of *PSPTO_3576* (*tvrR*) was not altered in the *pmeR* mutant (Fig 3D). Furthermore, we observed no significant differences in the auxin responsiveness of two auxin-downregulated genes, *PSPTO_1404* (*hrpL*) and *PSPTO_0371* (*iaaL*), between WT and the *pmeR* mutant (Figs 3E and 3F). These results indicate that *pmeR* not only locally regulates the auxin induction of *pmeR* but also modulates auxin responsiveness of several additional genes. Notably, the mutation of *pmeR* resulted in only a partial reduction in IAA induction of *PSPTO_1824*, *PSPTO_4297* and *aefR*, suggesting the involvement of at least one additional positive regulator in their auxin responsiveness.

**Fig 3.**
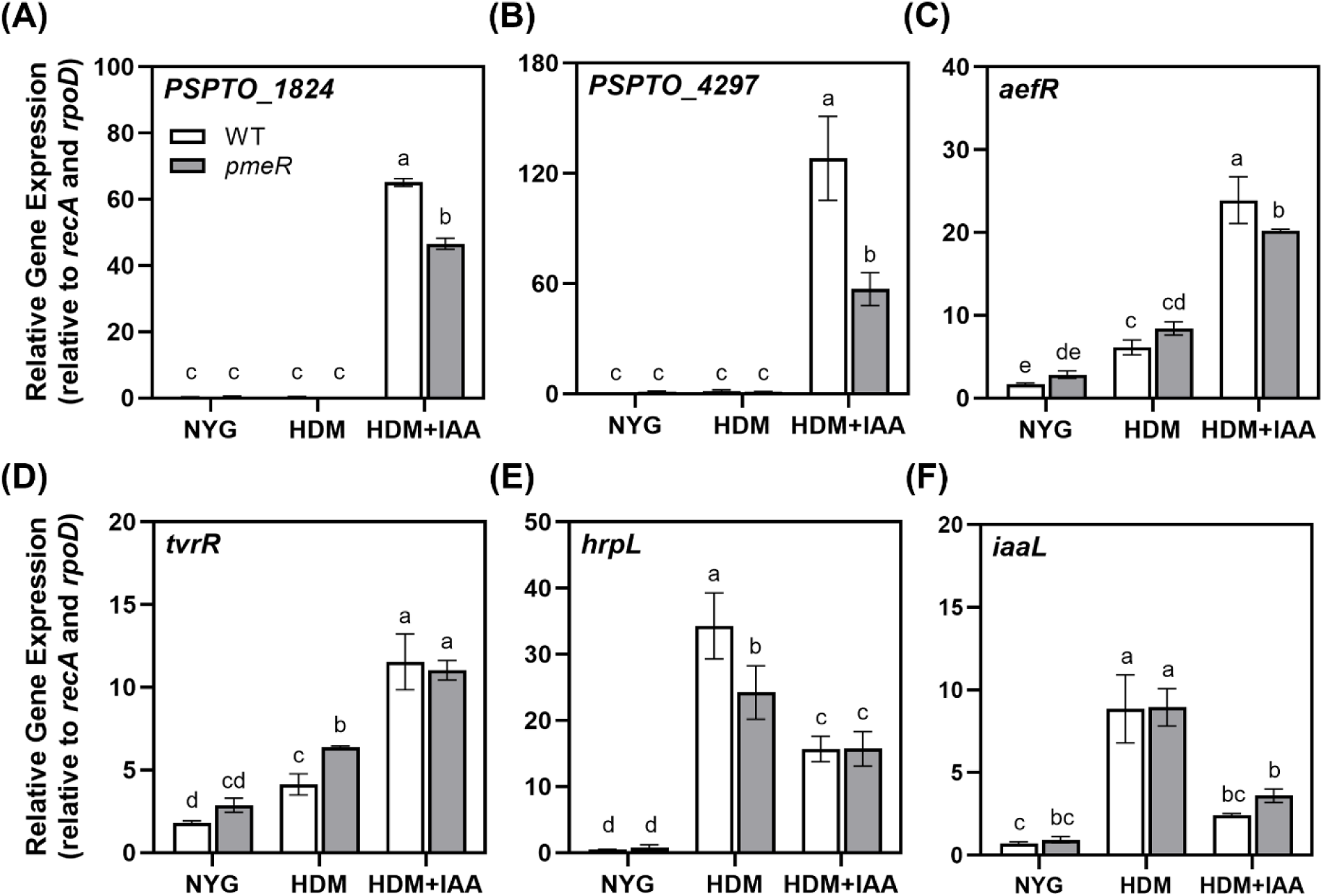
Mutation of *pmeR* alters *Pto*DC3000 auxin responsiveness in culture. Expression of auxin-responsive genes (A) *PSPTO_1824*, (B) *PSPTO_4297*, (C) *PSPTO_3549* (*aefR*), (D) *PSPTO_3576* (*tvrR*), (E) *PSPTO_1404* (*hrpL*), and (F) *PSPTO_0371* (*iaaL*) was quantified by qRT-PCR. Auxin responsiveness of the indicated genes in WT or *pmeR*::ΩKan mutant (*pmeR*) cells was monitored 30 minutes after transfer from NYG to the indicated medium: NYG, HDM, or HDM + IAA. Relative gene expression for each gene was calculated by normalizing to the expression of two reference genes, *rpoD* and *recA.* Data is from a representative experiment (n = 3) and plotted as the mean of relative gene expression ± SD. Similar results were obtained in a second independent experiment. Letters indicate significant differences between treatments as determined by ANOVA followed by Tukey’s HSD test (*p* < 0.05). NYG: nutrient yeast glycerol medium; HDM: Hrp de-repressing medium; HDM + IAA: HDM containing 100 µM IAA.

To confirm the role of *pmeR* in regulating auxin responsiveness in *Pto*DC3000, we generated an independent *pmeR* deletion mutant strain (*pmeR*::Δ, see Materials and Methods). We introduced the *pmeR_prom_-lux* reporter into *pmeR*::Δ, and observed de-repressed expression of *pmeR_prom_-lux*, confirming the functional disruption of *pmeR* in this strain (S2A Fig). We next monitored the auxin responsiveness of *PSPTO_4297* in the *pmeR*::Δ mutant by taking advantage of a previously described *PSPTO_4297_prom_-lux* transcriptional fusion (15). Consistent with our in-culture qRT-PCR results (Fig 3B), IAA strongly induced the expression of *PSPTO_4297_prom_-lux* in the wild-type strain, and the level of IAA induction of *PSPTO_4297_prom_-lux* was significantly reduced in the *pmeR*::Δ mutant (S2B Fig). These findings confirm that *pmeR* is required for normal auxin responsiveness of *Pto*DC3000 in culture.

The auxin-responsive genes discussed above have been shown to be upregulated during *Pto*DC3000 growth *in planta* (10, 17, 18). To investigate whether *pmeR* regulates these genes during pathogenesis, we analyzed their expression in *Pto*DC3000-infected *A. thaliana* leaves. We infiltrated *A. thaliana* with either WT or the *pmeR*::ΩKan mutant strain, and isolated total RNA from infected leaves at 6 hours post-infection (hpi) for qRT-PCR analysis, as previously described (9). Consistent with previous findings (10), we observed that in WT cells, expression of *pmeR* and all five of the selected auxin-responsive genes was induced at 6 hpi (S3 Fig and Fig 4). A comparison of transcript levels between WT and the *pmeR* mutant revealed significant differences in the expression patterns of all five of these bacterial genes *in planta*. Specifically, the induction levels of *PSPTO_1824*, *PSPTO_4297*, and *hrpL* were significantly lower in the *pmeR* mutant compared to WT (Figs 4A-4C), while the expression of *mexA* and *iaaL* was further augmented in the *pmeR* mutant (Figs 4D and 4E). These findings indicate that *pmeR* plays a crucial role in regulating the expression of these genes *in planta* and thus may contribute to the virulence of *Pto*DC3000.

**Fig 4.**
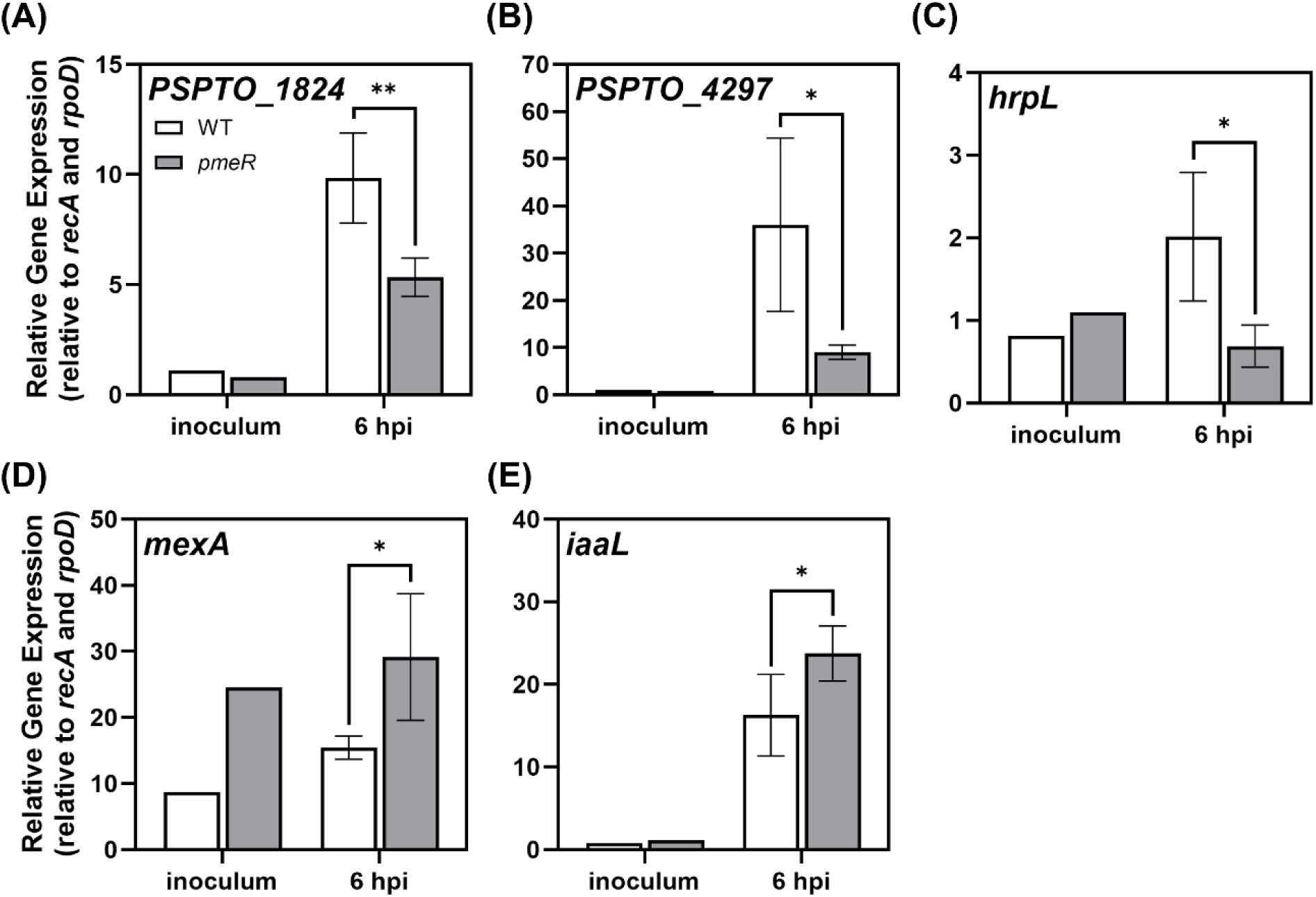
Mutation of *pmeR* alters expression of auxin-responsive *Pto*DC3000 genes *in planta*. Expression of bacterial genes (A) *PSPTO_1824*, (B) *PSPTO_4297*, (C) *PSPTO_1404* (*hrpL*), (D) *PSPTO_4303* (*mexA)*, and (E) *PSPTO_0371* (*iaaL*) in wild-type *Pto*DC3000 (WT) and the *pmeR*::ΩKan mutant (*pmeR*) growing in *A. thaliana* Col-0 plants. To quantify bacterial gene expression, infected leaves were harvested at 6 hours after inoculation, and total RNA was isolated and used for qRT-PCR. RNA isolated from the *Pto*DC3000 cell suspension used for inoculation (inoculum, n = 1) was used as the calibrator for relative expression. The relative expression for each gene was calculated by normalizing to the expression of two reference genes, *rpoD* and *recA.* Data is from a representative experiment (n = 3) and plotted as the mean of relative gene expression ± SD. Similar results were obtained in a second independent experiment. Asterisks indicate significant differences between WT and *pmeR* strains as determined by Student’s *t*-test (*: *p* < 0.05; **: *p* < 0.01).

### *pmeR* contributes to virulence of *Pto*DC3000 on *A. thaliana*

Given the positive role of auxin in enhancing *Pto*DC3000 virulence in *A. thaliana* (19) and that *pmeR* plays a role in regulating expression of auxin-responsive genes *in planta* (Fig 4), we hypothesized that the misregulation of auxin responsiveness in the *pmeR* mutant might impact its virulence. We first confirmed that the *pmeR*::ΩKan mutant had no growth defect in culture (S4 Fig). We next tested if the misregulation of auxin responsiveness in the *pmeR* mutant impacts its growth *in planta*. We infected *A. thaliana* with either the WT strain or the *pmeR*::ΩKan mutant via surface inoculation, tracking bacterial growth during pathogenesis. The growth of the *pmeR* mutant was significantly reduced at both 2- and 4-days post-inoculation (dpi) compared to the WT strain (Fig 5A). We also observed that leaves inoculated with the *pmeR* mutant developed fewer water-soaked lesions and less chlorosis at 4 dpi (Fig 5B). To determine if this compromised virulence was due to the *pmeR* mutant being impaired in entry into the leaf, we quantified the number of bacteria present inside leaves at 4 hours post-dip-inoculation. We observed no significant differences in bacterial number between the WT strain and the *pmeR* mutant within surface-sterilized leaves (Fig 5C). Additionally, we infected *A. thaliana* with either WT or the *pmeR*::ΩKan mutant using syringe inoculation to bypass the entry stage, and observed that the *pmeR* mutant exhibited significantly reduced growth at 2 and 3 dpi (Fig 5D). Thus, the *pmeR* mutant is impaired in the ability to grow in the leaf apoplast. To verify that the reduced virulence of the *pmeR* mutant was due to disruption of *pmeR* function, we tested if we could restore the virulence of the *pmeR*::ΩKan mutant by introducing the wild-type *pmeR* gene including its upstream regulatory region on a stable plasmid (*pmeR/*pME6031::*pmeR^+^*). As controls, this plasmid was also introduced into WT (WT*/*pME6031::*pmeR^+^*), and the empty vector (pME6031) was introduced into both WT (WT/pME6031) and the *pmeR*::ΩKan mutant (*pmeR*/pME6031). Consistent with our earlier infection assays (Fig 5), the *pmeR*/pME6031 strain exhibited significantly reduced bacterial growth at 2 and 4 dpi compared to the WT/pME6031 strain (S5 Fig). Growth of the *pmeR/*pME6031::*pmeR^+^* strain was consistently higher than that of the *pmeR*/pME6031 strain at 4 dpi, although the increase was not statistically significant. These findings demonstrate that *pmeR* is important for *Pto*DC3000 growth in the leaf apoplast, contributing to its virulence.

**Fig 5.**
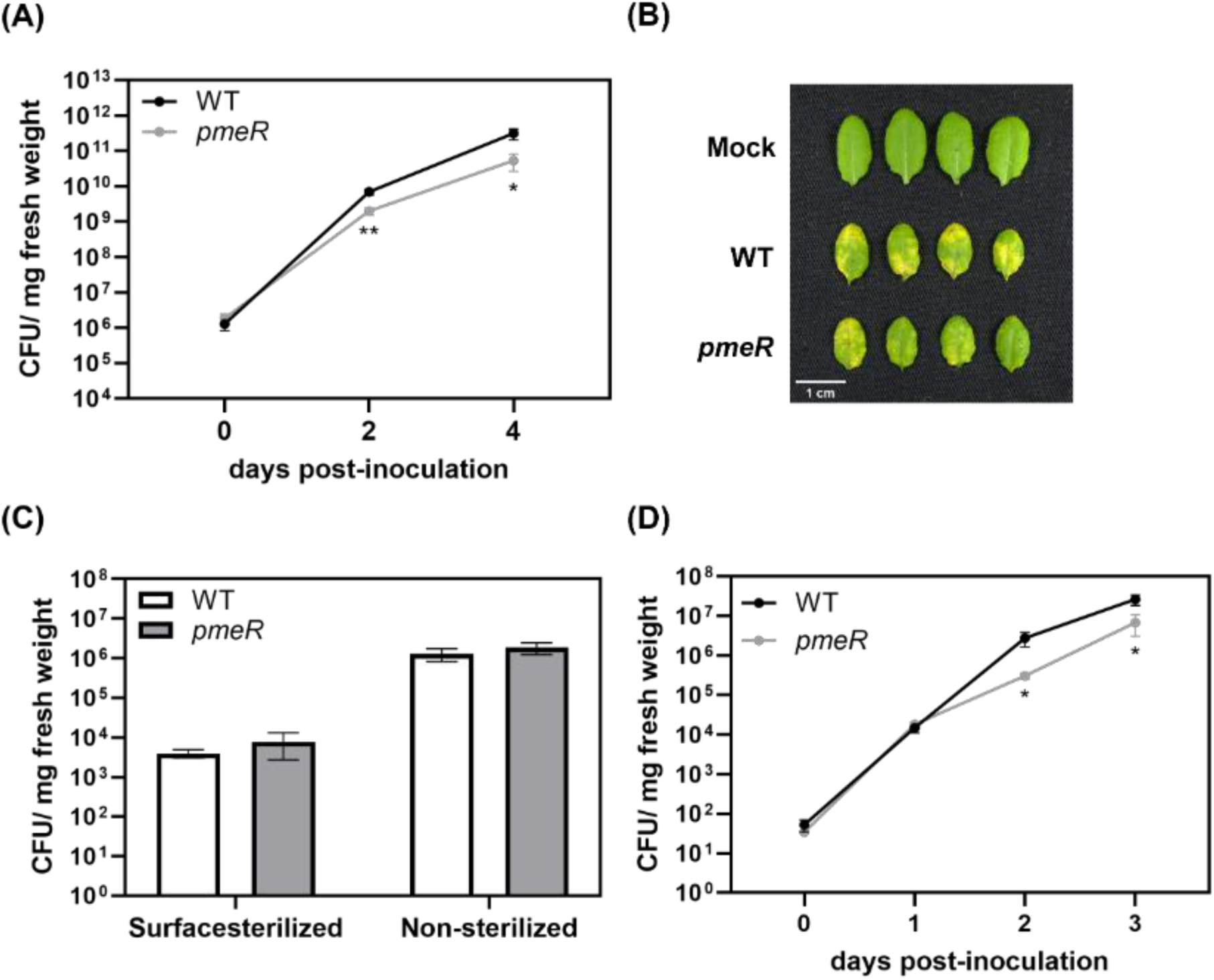
*pmeR* is required for full virulence of *Pto*DC3000 on *A. thaliana*. (A) Growth of WT and *pmeR*::ΩKan mutant (*pmeR*) bacteria in four-week-old *A. thaliana* Col-0. Plants were dip-inoculated with ∼ 5 x 10^7^ CFU/mL of bacteria. The total number of bacteria in infected leaves was quantified at 0, 2, and 4 dpi. Data from two independent experiments are combined and shown as mean ± standard error (SE) (n = 12 for 0 dpi, and n = 24 for 2 and 4 dpi). (B) Disease symptoms on dip-inoculated *A. thaliana* leaves at 4 dpi. Mock: Leaves dipped with 10 mM MgCl_2._ (C) Quantification of bacterial growth in and/or on leaves at 4 hours post-inoculation. Surface-sterilized leaves were treated with 15% hydrogen peroxide to remove bacteria from the surface of dip-inoculated leaves. Data are combined from two independent experiments and shown as mean ± SE (n = 8). (D) Growth of WT and *pmeR* mutant strains in four-week-old wild-type *A. thaliana* Col-0. Plants were infiltrated with ∼ 1 x 10^5^ CFU/mL of bacteria. The number of bacteria in infiltrated leaves was quantified at 0, 1, 2, and 3 dpi. Data from two independent experiments are combined and shown as mean ± SE (n = 12 for 0 dpi, and n = 24 for 1 to 3 dpi). Asterisks indicate significant differences between WT and *pmeR* mutant strains as determined by Student’s *t*-test (*: *p* < 0.05; **: *p* < 0.01). CFU: Colony forming units.

### PmeR does not appear to be a direct sensor of IAA

Upregulation of genes regulated by TFRs classically involves de-repression of transcription in response to the binding of small molecules, which are often substrates of the co-regulated efflux pumps encoded by adjacent operons. Based on our observations that IAA induces the expression of both *pmeR* and *mexA* (Fig 1) and a previous finding that at high concentrations (> 100 µM), IAA inhibits bacterial growth in culture (25), we hypothesized that IAA might be both a substrate for the MexAB-OprM efflux pump and a ligand for PmeR. To test if IAA is one of the substrates of the MexAB-OprM efflux pump, we performed minimal inhibitory concentration (MIC) assays to examine if the disruption of *mexA* decreases tolerance to IAA in *Pto*DC3000. In WT cells, 70 µg/mL (400 µM) of IAA completely inhibited its growth (Table 1). On the other hand, growth of the *mexA*::ΩKan mutant (*mexA*, (12)) was completely inhibited at 35 µg/mL (200 µM) of IAA. The reduced MIC observed in the *mexA* mutant is consistent with IAA being a substrate of MexAB-OprM efflux pump. We did not observe a difference in MICs between WT and the *pmeR* mutant (Table 1), presumably due to the fact that both IAA treatment of WT cells (Fig 1B) and the disruption of *pmeR* results in elevated expression of *mexA* (Fig 1B) (12, 20).

**Table 1.** The *mexA* mutant is more sensitive to IAA compared to WT.

| Compound ($\mu\text{g/mL}$) | WT | *pmeR* | *mexA* |
| --- | --- | --- | --- |
| IAA | $70 \pm 0$ | $70 \pm 0$ | $35 \pm 0$ |

The MICs of indicated compounds are shown as mean ± SD ( $\mu\text{g/mL}$ ) and are compiled from 3 independent experiments, consisting of 2 biological replicates each ( $n = 6$ ). WT: wild-type *PtoDC3000*; *pmeR*: *pmeR::ΩKan* mutant ; *mexA*: *mexA::ΩKan* mutant.

We next tested the hypothesis that IAA might be a ligand for PmeR, and that binding of IAA to PmeR would decrease the affinity of PmeR for its regulatory sequences and thus de-repress *pmeR* transcription. To test this, we carried out electrophoresis mobility shift assays (EMSAs) with purified PmeR protein and a DNA probe containing the intergenic region between *pmeR* and the *mexAB*-*oprM* operon (Fig 6A) that contains two previously identified overlapping palindromic PmeR binding sites (Fig 6A, (12)). Incubating the [γ-32P]-labeled DNA probe (1.5 nM) with increasing concentrations of purified PmeR protein (ranging from 0.39 to 25 nM) showed PmeR binding to the DNA in a concentration-dependent manner, with 6.25 nM being the concentration that resulted in a complete shift of the unbound DNA probe to the PmeR/DNA complex (Fig 6B). It is worth noting that we observed two sizes of shifted PmeR protein/DNA complexes in these experiments, presumably corresponding respectively to single and double occupancy of the two palindromic PmeR-binding sites in the regulatory region (Fig 6A). Competition assays showed that excess unlabeled DNA probe competitively inhibited the binding of PmeR to the labeled DNA probe (Fig 6B, lane 9), validating that PmeR specifically binds to the intergenic region between *pmeR* and the *mexAB*-*oprM* operon.

**Fig 6.**
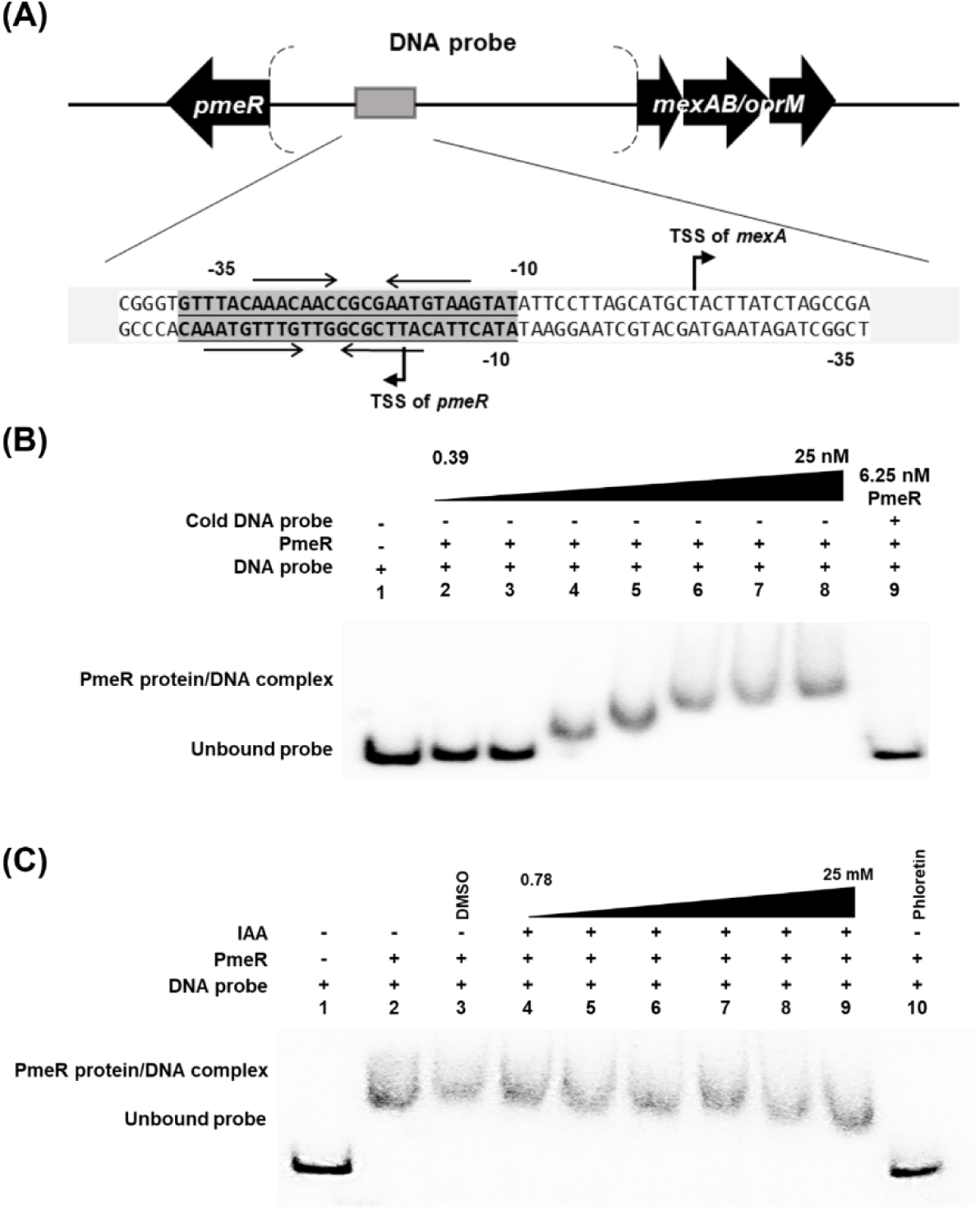
IAA does not impact the binding of PmeR to its regulatory region *in vitro*. (A) A schematic diagram (not drawn to scale) of the region of the *Pto*DC3000 genome encoding *pmeR* and *mexA*. The 254-bp intergenic region between the *Pto*DC3000 *pmeR* gene and the *mexAB*-*oprM* operon was used as the DNA probe in the EMSA shown in panels B and C. The nucleotide sequence of the regulatory region, the PmeR binding site (sequences in the grey box), and transcription start sites (TSS) for *mexA* and *pmeR* mapped by Vargas *et al*. (2011) (12) are shown. Horizontal arrows represent the nucleotides that interact with PmeR. (B) *In vitro* binding of purified PmeR protein to the *pmeR-mexA* intergenic region. Phosphor image of an EMSA using the [γ-32P]-labeled DNA probe amplified from the *pmeR* intergenic region. The probe (1.5 nM) was incubated with increasing concentrations (0.39 to 25 nM) of purified PmeR protein (Lanes 2 to 8). Lane 1 serves as a negative control in which only 1.5 nM DNA probe was added to the reaction buffer. Competitive EMSA was carried by the co-incubation of 6.25 nM PmeR with 1.5 nM labeled DNA probe and 30 nM unlabeled DNA probe (Lane 9). (C) Increasing concentrations of IAA (0.78 to 25 mM) were added to the co-incubation of 1.5 nM DNA probe and 6.25 nM PmeR (Lane 4 to 9). Lane 1 serves as a negative control to which only 1.5 nM DNA probe was added to the reaction buffer. Lane 2 presents the binding of PmeR to the DNA probe without chemical addition. The addition of DMSO solvent does not disrupt PmeR/DNA interaction (Lane 3), and the addition of 1 mM phloretin disrupts the PmeR/DNA interaction (Lane 10). Similar results were obtained in two additional independent experiments.

We then proceeded to use 6.25 nM of PmeR in further EMSAs to examine whether IAA impacts the binding of PmeR to the *pmeR* regulatory region. As a positive control, we included phloretin, a plant-derived flavonoid that has been reported to be a ligand of PmeR (12), in these assays and confirmed that 0.25 mM of phloretin caused the dissociation of PmeR from the DNA probe (S6A Fig). We co-incubated increasing concentrations of IAA (ranging from 0.78 to 25 mM) with 6.25 nM of PmeR and 1.5 nM of DNA probe. Although we observed that phloretin effectively caused the dissociation of PmeR from the DNA, binding of PmeR to the DNA probe was unaffected by the presence of IAA (Fig 6C), suggesting that IAA is not a ligand for PmeR. Thus, PmeR does not appear to be a direct intracellular IAA sensor. These findings prompted us to explore other potential signaling molecules that may regulate *pmeR*-dependent gene expression.

### What are the signals that regulate *pmeR*-dependent gene expression?

If IAA is not the ligand for PmeR, we hypothesized that IAA might potentially be converted into a related metabolite, such as 3-indole-acetyl-ɛ-L–lysine (IAA-Lys), upon entry into the *P. syringae* cell. *Pto*DC3000 does not encode any of the known IAA catabolism pathways and IAA-Lys is the only known IAA conjugate produced by *P. syringae* (21). Thus, we hypothesized that IAA-Lys could serve as a PmeR ligand, inducing gene expression in response to IAA.

To explore IAA-Lys and other potential IAA-related metabolites that might act as inducers of *pmeR*, we monitored the luminescence of *pmeR_prom_-lux* at 90 minutes post-treatment with IAA-Lys and two IAA-amino acid conjugates produced by plants: N-(3-Indolylacetyl)-L-alanine (IAA-Ala), and N-(indole-3-acetyl)-L-aspartic acid (IAA-Asp) (22). Among the IAA-amino acid conjugates tested, only IAA-Lys significantly induced expression of *pmeR_prom_-lux* in WT cells, although not as strongly as IAA (Fig 7A). This level of expression was similar to the de-repressed level observed in *pmeR* mutants in the absence of IAA. In the *pmeR* mutant cells, IAA-Lys did not further induce expression of *pmeR_prom_-lux* beyond the de-repressed levels, suggesting that the IAA-Lys induced expression may be fully dependent upon *pmeR*. For comparison, we also included phloretin and indole, as previous studies demonstrated that both molecules can de-repress *pmeR*-regulated gene expression (12, 20). Surprisingly, neither phloretin nor indole were effective inducers in our hands (Fig 7A). This could be due to the differences in growth media used in our studies. We conducted our reporter assays in HDM, which is a defined medium thought to mimic the acidic nutrient-limited conditions encountered in the apoplast, while the previous studies used rich media for their assays. Thus, neither of these molecules appeared to be a signal for the *pmeR-*dependent IAA responsiveness observed under our experimental conditions, suggesting that IAA-Lys could be a ligand of PmeR.

**Fig 7.**
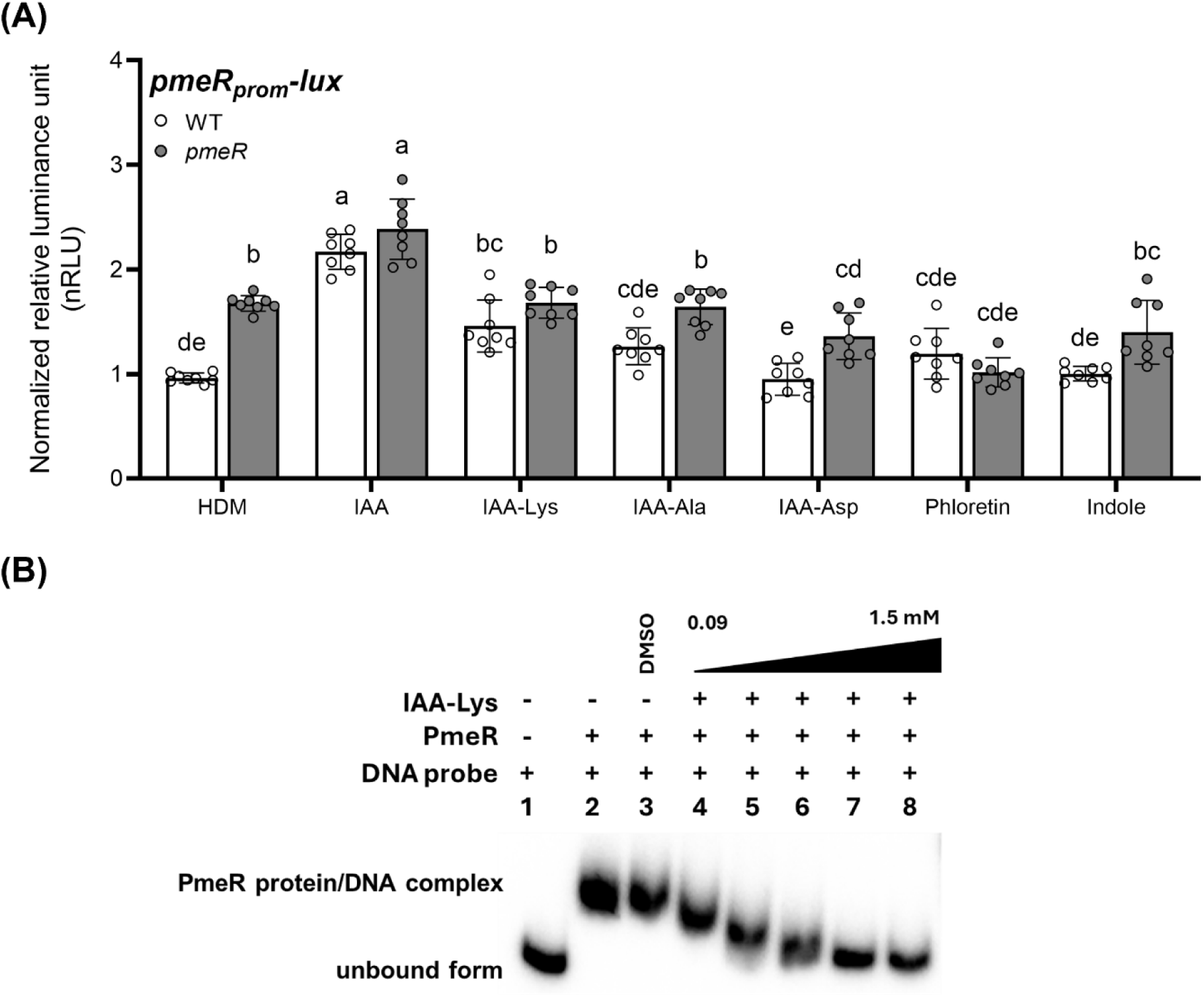
The auxin conjugate IAA-Lysine de-represses *pmeR* promoter activity. (A) Expression of the *pmeR_prom_-lux* reporter in WT and the *pmeR*::ΩKan mutant (*pmeR*) was quantified by monitoring luminescence in response to the indicated compounds (100 µM) at 90 minutes post-treatment. Data represent the mean of normalized relative luminescence (nRLU) ± SD. nRLU values were calculated by normalizing RLU to the mean value of *pmeR_prom_-lux* reporter activity in WT, in HDM. Data were compiled from two independent experiments (n = 8). Letters indicate significant differences between treatments as determined by ANOVA followed by Tukey’s HSD test (*p* < 0.05). HDM: Hrp de-repressing medium; IAA: indole-3-acetic acid; IAA-Lys: IAA-lysine conjugate; IAA-Ala: IAA-alanine conjugate; IAA-Asp: IAA-aspartate conjugate. (B) Increasing concentrations of IAA-Lys (ranging from 0.09 to 1.5 mM, Lane 4 to 8) were added to the co-incubation of 1.5 nM [γ-32P]-labeled DNA probe containing the intergenic region between *pmeR* and the *mexAB*-*oprM* operon and 6.25 nM PmeR. Lane 1 is a negative control that includes 1.5 nM DNA probe in the reaction buffer. Lane 2 shows the binding of the DNA probe and PmeR without chemical addition. Lane 3 serves as a control to show that the addition of DMSO solvent does not disrupt PmeR/DNA interaction. Similar results were obtained in two additional independent experiments.

To investigate whether IAA-Lys is a ligand of PmeR, we examined its ability to impact the binding of PmeR to its DNA binding site in our EMSA assay. We observed that co-incubation with increasing concentrations of IAA-Lys affected the binding of PmeR to the DNA probe (Fig 7B). Specifically, 750 µM of IAA-Lys was sufficient to completely shift the PmeR/DNA complex to the unbound DNA form (Fig 7B). These results suggest that IAA-Lys is a ligand for PmeR and plays a role in the auxin-induced expression of *pmeR*.

## Discussion

Upon infection with *Pto*DC3000, IAA accumulates within the infected leaf tissue, promoting pathogen growth and disease development (9, 19). One proposed mechanism by which IAA promotes *Pto*DC3000 virulence is through the reprogramming of bacterial gene expression (9, 10), and that IAA thus serves as a crucial signal for the transition between early and intermediate stages of *Pto*DC3000 pathogenesis (15). However, the mechanisms *Pto*DC3000 uses to sense and respond to IAA are unknown. In this study, we identified a novel bacterial auxin-signaling mechanism that is mediated by the auxin-upregulated transcription regulator, PmeR, and investigated its contribution to the virulence of *Pto*DC3000 on *A. thaliana*.

### PmeR regulates IAA-induced gene expression in *Pto*DC3000

Consistent with previous reports (12, 20), we demonstrated that PmeR functions as a transcriptional repressor that locally regulates the expression of its own gene and the adjacent *mexAB-oprM* operon (S1 Fig and Fig 4D). We also showed that both *pmeR* and *mexA* are auxin-upregulated genes (Fig 1), and that *pmeR* is required for *Pto*DC3000’s normal response to IAA in culture (Figs 2 and 3), underscoring its regulatory role in the *Pto*DC3000 auxin-sensing mechanism.

Given that several TFRs have been reported to directly regulate the transcription of multiple genes, not just those encoded locally (23, 24), we hypothesize that PmeR might more broadly regulate *Pto*DC3000 transcription in response to auxin. The exact mechanism by which PmeR induces expression of some auxin-responsive genes is unknown. It is possible that PmeR functions as a transcriptional activator that directly upregulates a subset of auxin-responsive genes (e.g., *PSPTO_1824*, *PSPTO_4297*, and *aefR*). In fact, although most TFRs act as either repressors or activators, some TFRs can function as both (11, 25), such as DhaS in *Lactococcus lactis* (26) and SMU.1349 in *Streptococcus mutans* (27). Alternatively, this *pmeR*-dependent regulation could be indirect, resulting from the de-repression of one or more unidentified *pmeR*-regulated genes that encode additional transcriptional regulators or via other molecular mechanisms. Further investigation using ChIP-seq followed by molecular genetics analyses will identify other PmeR- dependent regulons and elucidate their role in regulating *Pto*DC3000 auxin responsiveness.

Our findings indicate that *pmeR* is not the only regulator of the auxin response in *Pto*DC3000. For example, (i) IAA treatment further induced expression of *pmeR_prom_-lux* above the de-repressed level in both WT and the *pmeR* mutant at 90 minutes post-treatment (Fig 2) and (ii) the auxin induction of several genes (*PSPTO_1824*, *PSPTO_4297*, and *aefR*) was not completely abolished in the *pmeR* mutant (Figs 3 and 4). Thus, we hypothesize that at least one additional transcriptional regulator is involved in mediating auxin-responsive gene expression. It is possible that one more of the previously identified IAA-induced *Pto*DC3000 genes encoding known or putative transcriptional regulators or small RNAs could be involved in this process (10). Future investigations into the regulators governing auxin sensing and responses will further elucidate the mechanisms underlying bacterial auxin signaling in *Pto*DC3000.

### *pmeR* is required for full virulence of *Pto*DC3000 on *A. thaliana*

Our infection assays indicate that *pmeR* contributes to *Pto*DC3000 virulence on *A. thaliana*, as the *pmeR* mutant exhibits reduced growth in the apoplast (Fig 5). We propose that *pmeR*-mediated virulence is achieved through several different mechanisms. First, *pmeR* is likely to contribute to virulence through controlling the proper expression of auxin-responsive genes *in planta* (Fig 4). Some of these genes, such as *iaaL* (28), are known to contribute to *Pto*DC3000 virulence; while the roles of *PSPTO_1824* and *PSPTO_4297* in virulence remain unexplored. Second, *pmeR* could regulate virulence by ensuring the optimal expression of the *mexAB-oprM* operon during infection. We hypothesize that in addition to effluxing antimicrobial compounds, the MexAB-OprM pump may also regulate the intracellular concentration of signaling molecules (e.g. IAA, Table 1). For example, overexpression of the *mexAB-oprM* operon in the *pmeR* mutant might result in reduced intracellular levels of IAA or related molecules, which would impact the expression of auxin-responsive virulence genes. Third, we also observed that the disruption of *pmeR* impairs induction of the *hrpL* gene, an important virulence regulator (29, 30), both in culture (in HDM) and *in planta* (Figs 3 and 4). Thus, *pmeR* may regulate the expression of *hrpL* in response to unidentified signals during infection.

### The auxin conjugate IAA-Lys is a possible ligand of PmeR

Although IAA was the strongest inducer of *pmeR* expression (Fig 7A), our EMSA showed that IAA had no effect on PmeR binding to the *pmeR* regulatory region (Fig 6C), suggesting that IAA is not a ligand of PmeR and thus that PmeR is not a direct sensor of IAA. Flavonoids (e.g. phloretin) were previously reported to induce *pmeR* expression and interfere with the binding between PmeR and its operator (12); however, in our hands, phloretin did not induce expression of *pmeR_prom_*-*lux* (Fig 7). Moreover, we did not find any evidence in the *Pto*DC3000 genome for genes involved in flavonoid biosynthesis. Thus, the observed IAA induction of *pmeR* expression is not likely to be due to the IAA-induced production of flavonoids in *Pto*DC3000.

We propose that upon entry into the *Pto*DC3000 cell, IAA is converted into IAA-Lys or a similar molecule. The binding of PmeR to this molecule results in de-repression of *pmeR* and other *pmeR*- dependent genes. The results from our transcriptional and biochemical experiments, which showed that IAA-Lys can both induce *pmeR* expression in culture (Fig 7A) and disrupt the PmeR/DNA interaction (Fig 7B), support this hypothesis. To date, IAA-Lys is the only known auxin-amino acid conjugate produced by *P. syringae*. Previous studies in *P. syringae* and *P. savastanoi* strains have shown it is synthesized through the enzymatic action of IAA-Lys synthase, encoded by the *iaaL* gene (28, 30–33). However, Pintado *et al*. found that the IAA-Lys synthase encoded by the *Pto*DC3000 *iaaL* gene is much less active than the enzymes of several other *P. syringae* strains (21). Further investigations are needed to elucidate the role of IAA-Lys in IAA signaling and virulence of *Pto*DC3000.

### Conclusions and Future Perspectives

Our results highlight the relevance of *pmeR* in regulating auxin-responsive genes and pathogenicity of *Pto*DC3000. As summarized in Fig 8, we propose that PmeR binds to the regulatory region of its own gene and other *pmeR*-regulated genes, repressing their expression early during infection when IAA levels are low. As the concentration of IAA in the infected leaf increases, IAA moves into the bacterial cell, reprogramming bacterial gene expression through either *pmeR*-dependent or *pmeR*-independent mechanisms. Given that IAA does not appear to be a ligand for PmeR, we further hypothesize that IAA induces the accumulation of IAA-Lys or a similar molecule, which acts as a ligand of PmeR. Upon ligand binding, a conformational change in the protein occurs, releasing PmeR from its DNA regulatory region, resulting in de-repression of *pmeR*-regulated genes. Our gene expression data indicate that at least one additional transcriptional regulator is involved in mediating auxin-responsive gene expression in *Pto*DC3000. This observation suggests that *Pto*DC3000 has evolved multiple mechanisms to respond to auxin, highlighting the importance of auxin responsiveness in this interaction. The *pmeR*-dependent expression of *mexAB-oprM* likely contributes directly to virulence, as the MexAB-OprM pump effluxes antimicrobial compounds, such as plant-derived flavonoids, thus promoting growth of *Pto*DC3000 in the plant (12). An additional mechanism for modulating IAA-dependent gene expression may occur via regulation of the MexAB-OprM efflux pump, as it may impact the intracellular levels of IAA or related signaling molecules. The *pmeR*-independent pathway responsible for induction of genes in response to IAA presumably involves auxin-responsive transcriptional regulators that have not yet been identified.

**Fig 8.**
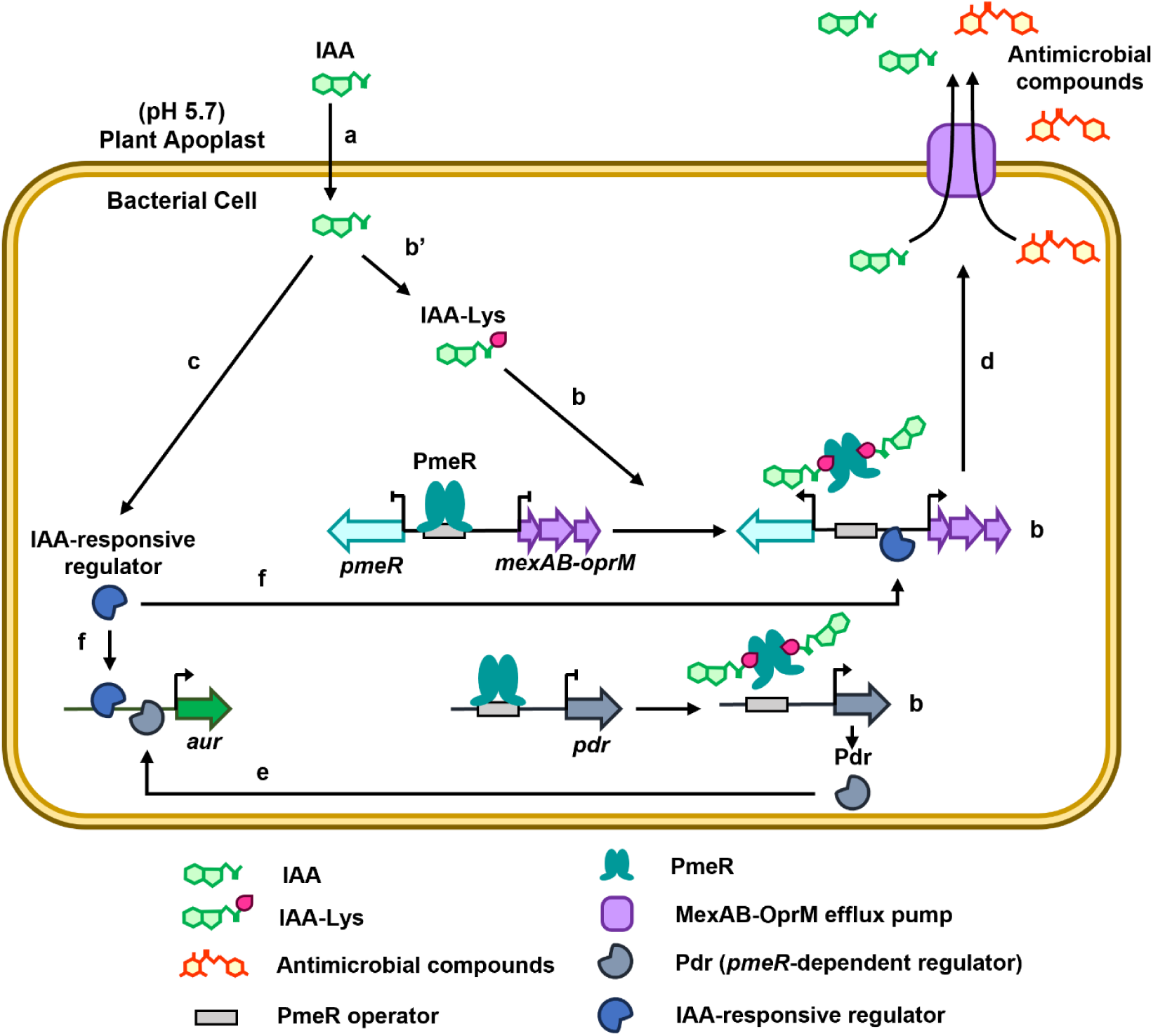
A working model for the role of PmeR in regulating IAA responses and virulence in *Pto*DC3000. *Pto*DC3000 infection leads to the accumulation of IAA within infected leaf tissue. Apoplastic IAA (which is largely in its protonated form at pH5.7 in the apoplast) can diffuse through the bacterial plasma membrane into the cell (a), where it reprograms bacterial gene expression through either *pmeR*-dependent (b) or *pmeR*-independent (c) regulatory mechanisms. The *pmeR* gene is upregulated by IAA, and encodes PmeR, a transcriptional regulator that, in the absence of its ligand, represses transcription of its target genes, *pmeR* and the adjacent *mexAB-oprM* operon. Binding of PmeR to its ligand decreases the affinity of PmeR for the operator region of the DNA, leading to the de-repression of both *pmeR* and *mexAB-oprM*. The IAA-induced transcription of *mexAB-oprM,* which encodes an efflux pump, leads to enhanced tolerance to plant-derived antimicrobial compounds, as well as to IAA itself (d). As the IAA-amino acid conjugate IAA-Lys (and not IAA) appears to be a ligand for PmeR, we propose that IAA is converted to IAA-Lys or a similar molecule upon entry into the cell (b’). We propose that PmeR also regulates the expression of one or more unidentified genes (*pdr*: *pmeR*-dependent regulator), which encode transcription factors that activate expression of auxin-upregulated (*aur*) genes (e) that are not direct targets of PmeR. The *pmeR*-independent pathway responsible for induction of genes in response to IAA presumably involves auxin-responsive transcriptional regulators that have not yet been identified (f). Ultimately, we propose that the upregulation of genes encoding efflux pumps and other virulence factors in response to IAA promotes the growth of *Pto*DC3000 in plant tissue.

To conclude, this study broadens the current understanding of bacterial auxin sensing mechanisms, and how complex regulatory networks facilitate infection of phytopathogenic microbes *in planta*. Future studies to identify additional regulators of auxin-responsive gene expression during pathogenesis will be valuable, as they will provide important insight into the molecular mechanisms underlying pathogen virulence strategies. Given the promising results with IAA-Lys as a ligand of PmeR, future studies to explore IAA-Lys as a potential signaling molecule will further elucidate mechanisms used by bacteria to sense and respond to IAA. Moreover, considering that several auxin-responsive genes that could contribute to virulence are regulated by *pmeR*, a thorough characterization of these genes would extend our knowledge of how auxin-responsive virulence factors contribute to the interaction between plant-associated microbes and their hosts.

## Materials and Methods

### Bacterial strains, plasmids, and culture conditions

The bacterial strains and plasmids used in this study are summarized in S1 Table. The *pmeR*::ΩKan and *mexA*::ΩKan mutants have been previously described (12). *P. syringae* wild-type *Pto*DC3000 (34) and mutant strains were grown in rich nutrient yeast glycerol (NYG) medium (35) or in defined hrp-derepressing medium (HDM) (36) supplemented with citrate (20 µM) and fructose (10 mM) at 28 ℃. *Escherichia coli* (*E. coli*) was grown on Luria-Bertani broth (LB) medium (37) at 37 ℃. Antibiotics used for selection included rifampicin (Rif, 80 µg/mL), kanamycin (Kan, 25 µg/mL), tetracycline (Tet, 16 µg/mL for *P. syringae* and 10 µg/mL for *E. coli*), spectinomycin (Spec, 100 µg/mL), chloramphenicol (Cm, 20 µg/mL), and amphotericin B (Amph, 5 µg/mL).

To monitor the growth of bacterial strains in culture, cells were grown overnight in NYG pH 7.0, sub-cultured (1:10 dilution) in NYG pH 7.0, and grown for 3 hours with aeration to mid-log phase. Cells were harvested by centrifugation, resuspended in 10 mM MgCl_2_ to an OD_600_ of 1.0, and 1 µL of the resuspension was inoculated into 100 µL of indicated media in sterile 96-well plates. The plates were incubated at 28 ℃ with shaking for 22 hours in an Epoch 2 microplate reader (Bio Tek), monitoring OD_600_ every hour.

### Plant material and growth conditions

All *A. thaliana* plants used in this study were in the Col-0 background. Plants were grown on soil in a nursery chamber with a short-day photoperiod (8 hours light/16 hours dark) at 21 ℃ and 50 % relative humidity, with a light intensity of ∼ 175 μEinsteins sec^-1^ m^-1^. Before inoculation experiments, the plants were moved to an infection chamber with similar growth conditions, but with higher relative humidity (75 %).

### *P. syringae* inoculation and quantification of bacterial growth

To monitor bacterial growth in planta, four- to five-week-old *A. thaliana* Col-0 (WT) plants were inoculated with the indicated *P. syringae* strains using either infiltration or surface inoculation as described previously (8, 9). In infiltration experiments, bacterial suspensions of 10^5^ cells/mL in 10 mM MgCl_2_ were introduced into fully expanded leaves using a 1-mL needleless syringe. For bacterial growth quantification, whole leaves were sampled at 4 hours after inoculation (day 0) (n = 4), and 2 and 4 days-post-inoculation (dpi) (n = 12), unless otherwise noted. Each inoculated leaf was weighed and then homogenized in 0.2 mL 10 mM MgCl_2_, using a MM 300 Laboratory Mixer Mill (Retsch). The resulting leaf extracts were serial diluted and plated onto NYG containing Rif (day 0 samples were plated onto NYG containing Rif and Amph to reduce fungal contamination). Following 48 hours of incubation at 28 ℃, the colonies were counted, and the number of colony forming units per milligram of leaf fresh weight (CFU/mg FW) was calculated.

Surface inoculation was carried out by dipping entire leaf rosettes of *A. thaliana* in bacterial suspensions of 5 x 10^7^ cells/mL in 10 mM MgCl_2_ containing 0.02 % Silwet L-77 surfactant (OSi Specialties). At four hours post-inoculation, fully expanded inoculated leaves were sampled. To evaluate the ability of *P. syringae* strains to enter into the leaf interior, surface-inoculated leaves were divided into two groups. One group of leaves was weighed and homogenized without surface sterilization, and the other leaves were surface sterilized with 15 % hydrogen peroxide (H_2_O_2_) for 5 minutes (38). After surface sterilization, the leaves were washed twice with sterile water, dried by dabbing with tissue to remove excess moisture, weighed and homogenized in 0.2 mL 10 mM MgCl_2_. The extraction solutions were then plated onto NYG containing Rif and Amph, and the number of colony forming units was counted after 48 hours incubation at 28 ℃. The quantification of bacterial growth at 2 and 4 dpi was determined by the same procedure described above. Disease symptoms were photographed at 4 dpi.

### Construction of *pmeR* promoter-luciferase (*lux*) reporter fusion and *pmeR* genomic clone

To construct the *pmeR_prom_*-luciferase reporter fusion, the regulatory region of *pmeR* (*PSPTO_4302*) was inserted upstream of the *luxCDABE* operon carried on pME6031::*lux* (29). Specifically, a fragment carrying the first 15 bp of the *pmeR* coding sequence and ∼ 250 bp of the upstream genomic sequence was amplified using primers FPlux KpnI-pmeR and RPlux EcoRI-pmeR (S2 Table). The resulting PCR product was digested with KpnI and EcoRI (Thermo Scientific) and ligated into the pME6031*::lux* plasmid digested with the same enzymes. The resulting plasmid was transformed into *E.coli* DH5α (39), sequenced to confirm its structure and that no mutations had been introduced during PCR, and then introduced into the indicated *Pto*DC3000 strains by triparental mating (40) and selection for Tet resistance.

The pME6031::*pmeR*^+^ plasmid, carrying the wild-type *pmeR* gene, was generated to complement the *pmeR*::ΩKan mutant. This was accomplished by amplifying a 936-bp DNA fragment including the 672-bp *pmeR* (*PSPTO_4302*) coding sequence, 254 bp of upstream and 10 bp of downstream genomic DNA, using primers FPlux KpnI-pmeR and FPlux KpnI-pmeR (S2 Table). The resulting DNA fragment was digested with KpnI and EcoRI (Thermo Scientific) and ligated into the pME6031 plasmid (14), which was digested with the same enzymes. The resulting plasmid was transformed into *E. coli* DH5ɑ (39), sequenced to confirm that no mutations had been introduced during PCR, and conjugated into the WT *Pto*DC3000 and *pmeR*::ΩKan strains by triparental mating (40) and selection for Tet resistance.

The helper plasmid pRK2013 (41) was used to mobilize the plasmids described above from *E.coli* DH5α (39) into *Pto*DC3000 WT and mutant strains (S1 Table).

### Construction of *pmeR* deletion mutant strain

To construct a second independent *pmeR* deletion mutant strain, a *pmeR* deletion plasmid was constructed in pK18mobsacB, a plasmid that allows for sucrose counterselection (42, 43). Approximately 1000 bp long genomic fragments flanking the *Pto*DC3000 *pmeR* (*PSPTO_4302*) coding region were amplified using Q5^®^ High-Fidelity DNA Polymerase (New England Biolabs) and the primers listed in S2 Table using the Gibson Assembly cloning method. The pmeRdownstreamFWD and pmeRupstreamREV primers were designed to result in an in-frame deletion in which only the first four and last four codons of the *pmeR* coding sequence are present (S2 Table). The amplified DNA fragments were assembled using the NEBuilder HiFi DNA Assembly^®^ Cloning Kit (New England Biolabs) following the manufacturer’s instructions. The resulting pK18*pmeR*Δ plasmid was transformed into *E.coli* DH5ɑ (39), sequenced to verify its structure and sequence, and conjugated into wild-type *Pto*DC3000 by triparental mating (40), and selecting for Rif and Kan resistance. The helper plasmid pRK2013 (41) was used to mobilize pK18*pmeR*Δ from *E.coli* DH5α (39) into *Pto*DC3000 (S1 Table).

Several Kan-resistant *Pto*DC3000 strains carrying the pK18*pmeR*Δ plasmid integrated at the *pmeR* locus were subjected to sucrose counterselection to screen for strains carrying the deletion of *pmeR*. Kan-resistant strains were plated onto NYG plates containing Rif and 15 % (v/v) sucrose to enrich for strains in which the *sacB* gene was lost, either through a double-crossover homologous recombination event (replacing the WT *pmeR* gene with the *pmeR*::Δ deletion allele), or a cross-over that looped out the entire plasmid, reverting the chromosome back to the original WT sequence. After a 4-day incubation at 28 ℃, sucrose-resistant colonies were patched onto NYG plates containing either Rif alone or both Rif and Kan. Kan-sensitive colonies were purified, and genotyped via colony PCR to identify strains in which the *pmeR*::Δ deletion had replaced the WT *pmeR* gene. A Kan-sensitive revertant strain was also purified to use as a control in future experiments.

### Bioluminescence assays to monitor *lux* reporter expression

To quantify bioluminescence, overnight bacterial cultures grown in NYG with the appropriate antibiotics were diluted 1:10 in NYG lacking antibiotics and grown for 3 hours at 28 ℃. When the cell density (OD_600_) of the subcultures reached approximately 0.2, the cells were collected by centrifugation and resuspended at an OD_600_ of approximately 2.0 in 10 mM MgCl_2_. The resuspensions were used to inoculate replicate cultures of 200 µL of HDM supplemented with 100 µM of the indicated compounds or with 0.1 % DMSO (as a solvent control) in a clear 96-well microplate (Greiner Bio-One) to an OD_600_ of approximately 0.025 - 0.10. The following compounds were tested: IAA (GoldBio), IAA-Lys (WuXi AppTec), IAA-Ala (Sigma-Aldrich), IAA-Asp (Sigma-Aldrich), phloretin (Sigma-Aldrich), and indole (Sigma-Aldrich). At indicated timepoints (30, 60, and 90 minutes post-inoculation), 100 µL of each bacterial culture were transferred to a LUMITRAC™ 200 microplate (Greiner Bio-One), and bioluminescence monitored using an Infinite M200PRO microplate reader (Tecan) with 1-second integration for luminescence readings. The OD_600_ of the bacterial cultures was also recorded using an Infinite M200PRO microplate reader (Tecan). To calculate relative luminescence values (RLU), the bioluminescence values, after subtracting the blank values (media only), were normalized per OD_600_. For each experiment, a minimum of three biological replicates were grown for each strain and treatment.

### IAA susceptibility testing

Minimal inhibitory concentration (MIC) assays were used to determine the susceptibility of *P. syringae* to IAA. Bacterial cells were grown overnight in NYG pH 7.0, sub-cultured (1:10 dilution) in NYG pH 5.7, and grown for 3 hours with aeration to mid-log phase. Cells were collected by centrifugation, resuspended in NYG pH 5.7 to OD_600_ of 1.0, and 1 µL of resuspension was inoculated into a series of 100 µL cultures in sterile 96-well plates. These cultures consisted of NYG pH 5.7 containing a series of two-fold dilutions of IAA. The plates were incubated at 28 ℃ with shaking for 20 hours. After 20 hours of growth, the lowest concentration of IAA that prevented growth of the culture was determined using the OD_600_ readings.

### RNA and complementary DNA (cDNA) preparation

Bacterial RNA and cDNA were prepared for quantitative reverse transcriptase PCR (qRT-PCR) to monitor differential gene expression under different conditions, as previously described (9). To monitor *Pto*DC3000 auxin-responsive gene expression in culture, total RNA was extracted from bacterial cells grown in HDM in the presence or absence of 100 µM IAA. Specifically, triplicate subcultures of WT *Pto*DC3000 and the *pmeR*::ΩKan mutant were first grown for 3 hours in NYG at 28 ℃ until they reached an OD_600_ of ∼ 0.2. The cells were then collected by centrifugation and resuspended to an OD_600_ of approximately 2. A volume of 100 µL of the bacterial resuspension was used to inoculate triplicate cultures consisting of 2.5 mL of NYG, HDM supplemented with 0.1% DMSO or with 100 µM IAA (dissolved in 0.1% DMSO). At 30 minutes post-treatment, 1.2 mL of culture from each treatment was sampled and mixed with 2 mL of RNAprotect Bacteria Reagent (Qiagen), flash frozen in liquid nitrogen, and stored at - 80 ℃. Frozen cell samples were thawed at room temperature and enzymatically lysed by adding 100 μL of 1 mg/mL lysozyme in TE buffer (10 mM Tris-HCl containing 1 mM EDTA•Na_2_), and total RNA extracted using the RNeasy RNA isolation kit (Qiagen) followed by on-column RNase-free DNase I treatment (Qiagen) according to the manufacturer’s instructions.

To monitor *Pto*DC3000 gene expression *in planta*, leaves of four-week-old *A. thaliana* Col-0 plants were inoculated with 10^6^ cells/mL of either WT *Pto*DC3000 or the *pmeR*::ΩKan mutant resuspended in 10 mM MgCl_2_. Triplicate samples of approximately 100 mg of inoculated leaves were collected for total RNA isolation at 6 hours post-inoculation, flash-frozen in liquid nitrogen, and stored at −80°C. Total RNA was extracted following a combination of protocols from the RNAprotect bacterial reagent kit (Qiagen) and RNeasy plant minikit (Qiagen) as described by Djami-Tchatchou *et al.* (9). As a calibrator of relative gene expression *in planta*, bacterial RNA from 1 mL of the initial inoculum was isolated using the same RNA extraction protocol for in-culture samples described above.

The RNA samples were quantified using a NanoDrop™ One Spectrophotometer (Thermo Scientific), and flash frozen in liquid nitrogen for storage. Approximately 1 mg of total RNA was used for cDNA synthesis using the RevertAid First Strand cDNA Synthesis kit (Thermo Scientific) and random hexamers (Integrated DNA Technologies) as primers. Control reactions without the addition of reverse transcriptase were included as quality controls to ensure no genomic DNA contamination was present in the RNA samples.

### Bacterial gene expression analysis

qRT-PCR was used to monitor the transcript levels of genes of interest using Ig^TM^ SYBR Green qPCR 2X Master Mix (Intact Genomics) on a CFX Connect Real-Time PCR System (Bio-Rad) with the following cycling conditions: 15 minutes at 95 ℃ followed by 40 cycles of 95 ℃ for 5 seconds and 58 ℃ for 30 seconds. A melt curve was generated to determine the specificity of each amplicon after the 40^th^ cycle, using these parameters: 65 °C for 5 sec, 95 °C for 5 min, then a slow ramp (0.5 °C for 5 sec), with camera capture. In each experiment, gene expression analysis was performed on three biological replicates with two technical replicates for each. The cycle threshold (Ct) of each gene was normalized to *rpoD* (*PSPTO_0537*) and *recA* (*PSPTO_4033*), which were used as reference genes as described in previous bacterial gene expression studies for growing *in planta* (9, 44). All primers used for qRT-PCR are described in S2 Table.

### Protein expression and purification

To generate a plasmid expressing an N-terminal His-tagged PmeR protein for use in electrophoretic mobility shift assay (EMSA), the coding region of *Pto*DC3000 *pmeR* was codon-optimized for expression in *E. coli*, synthesized, and ligated into pET-28a-c (+) vector by Twist Bioscience (South San Francisco, CA), and transformed into *E. coli* BL21 (DE3) cells (S1 Table). An overnight culture of transformed *E. coli* cells grown in LB containing 25 µg/mL Kan was diluted 1:100 dilution into 500 mL LB media containing 25 µg/mL Kan and grown at 37 ℃. When the culture had reached an OD_600_ of approximately 0.3, the culture was cooled down to 16 ℃ in a refrigerated shaker and induced with 1 mM isopropyl 1-thio-β-D-galactopyranoside (IPTG) at 16 ℃ with aeration for 20 hours. The induced cells were then harvested by centrifugation at 4,000 rpm at 4 ℃ for 30 minutes and stored at −80 ℃.

To purify the PmeR protein, we followed a previously described protocol for purification of a related TFR (20), with the following modifications. To purify the His-tagged PmeR protein, the cell pellets were resuspended in lysis buffer containing 20 mM Tris-HCl (pH 8.0), 20 mM NaCl, 5 mM imidazole, 5 % (vol/vol) glycerol, and 1mM phenylmethylsulfonyl fluoride (PMSF), and sonicated on ice to lyse the cells. The lysed culture was then centrifuged at 15,000 rpm at 4 ℃ for 45 minutes, and the supernatant passed over a nickel column equilibrated with lysis buffer lacking PMSF. The column was washed with 10-fold column volume of wash buffer containing 20 mM Tris-HCl (pH 8.0), 20 mM NaCl, 10 mM imidazole, and 5 % (vol/vol) glycerol. The bound His-tagged protein was eluted with elution buffer containing 20 mM Tris-HCl (pH 8.0), 20 mM NaCl, 300 mM imidazole, and 5 % (vol/vol) glycerol, and dialyzed overnight with elution buffer lacking imidazole at 4 ℃. After dialysis, samples were aliquoted and stored in −80 ℃. Protein concentration was determined using the Bradford method with BSA as a standard.

### Electrophoretic mobility shift assay (EMSA)

A 269-bp DNA probe containing the entire 254-bp *pmeR-mexA* intergenic region and 15-bp of the *pmeR* coding sequence was PCR-amplified from *Pto*DC3000 chromosomal DNA using Q5^®^ High-Fidelity DNA Polymerase (New England Biolabs) and primers FPlux KpnI-pmeR and RPlux EcoRI-pmeR (S2 Table). The resulting PCR product was purified using PureLink^TM^ PCR purification kit (Invitrogen) and radiolabeled at the 5’-hydroxyl termini with [γ-32P] ATP (Revvity) and T4 polynucleotide kinase (New England Biolabs).

The labeled probe was incubated with the indicated concentrations of purified His-tagged PmeR protein and chemicals of interest in 10 µL of modified STAD buffer (25 mM Tris-acetate [pH 8.0], 8 mM Mg-acetate, 10 mM KCl, 1 mM DTT, and 460 µM HCl) (12). 100 µg/mL of BSA (Thermo Scientific) and 25 µg/mL of poly dI-dC (Sigma-Aldrich) were added to the reaction to prevent non-specific interaction between PmeR protein and labeled DNA probe. Selected compounds at indicated concentrations were added to reactions: phloretin (Sigma-Aldrich), IAA (GoldBio), indole (Sigma-Aldrich), and IAA-Lys (WuXi AppTec). The reaction was incubated at 25 ℃ for 30 minutes and run on 4 % Tris-Borate (TB) native polyacrylamide gel for 1.5 hours at 80 V in pre-chilled TB buffer. The radioactively labeled probes were imaged and analyzed using an Amersham™ Typhoon™ IP (Cytiva).

### Statistical analysis

Datasets were statistically compared with Microsoft Excel and the statistical analysis software R 1.3.1073 (R Core Team, 2021). Statistical tests were performed using either Student’s *t*-test or one-way analysis of variation (ANOVA) followed by Tukey’s HSD test when appropriate. The confidence level of all analyses was set at 95 %, and values with *p* < 0.05 were considered significant.

## Acknowledgments

We thank Dr. Joshua M. B. Johnson for constructing the *PSPTO_4297_prom_-lux* reporter and Dr. María-Trinidad Gallegos for generously providing the *Pto*DC3000 *pmeR*::ΩKan and *mexA*::ΩKan insertion mutants. We thank Stefanie F. King for performing the initial characterization of the auxin responsiveness of the *pmeR*::ΩKan mutant and for helpful comments on the manuscript. We thank Dr. Hani S. Zaher, Dr. Sarah D. Beagle, and Dr. Jae S. Morris for their valuable assistance with EMSA, MIC, and protein purification experiments, respectively. We thank the staff of the Jeanette Goldfarb Plant Growth Facility for providing plant growth facilities and care for the plants. Finally, we thank Dr. Arnaud T. Djami-Tchatchou and Dr. Joshua M. B. Johnson for their helpful comments on the manuscript.

## Supporting information captions

**Fig S1.**
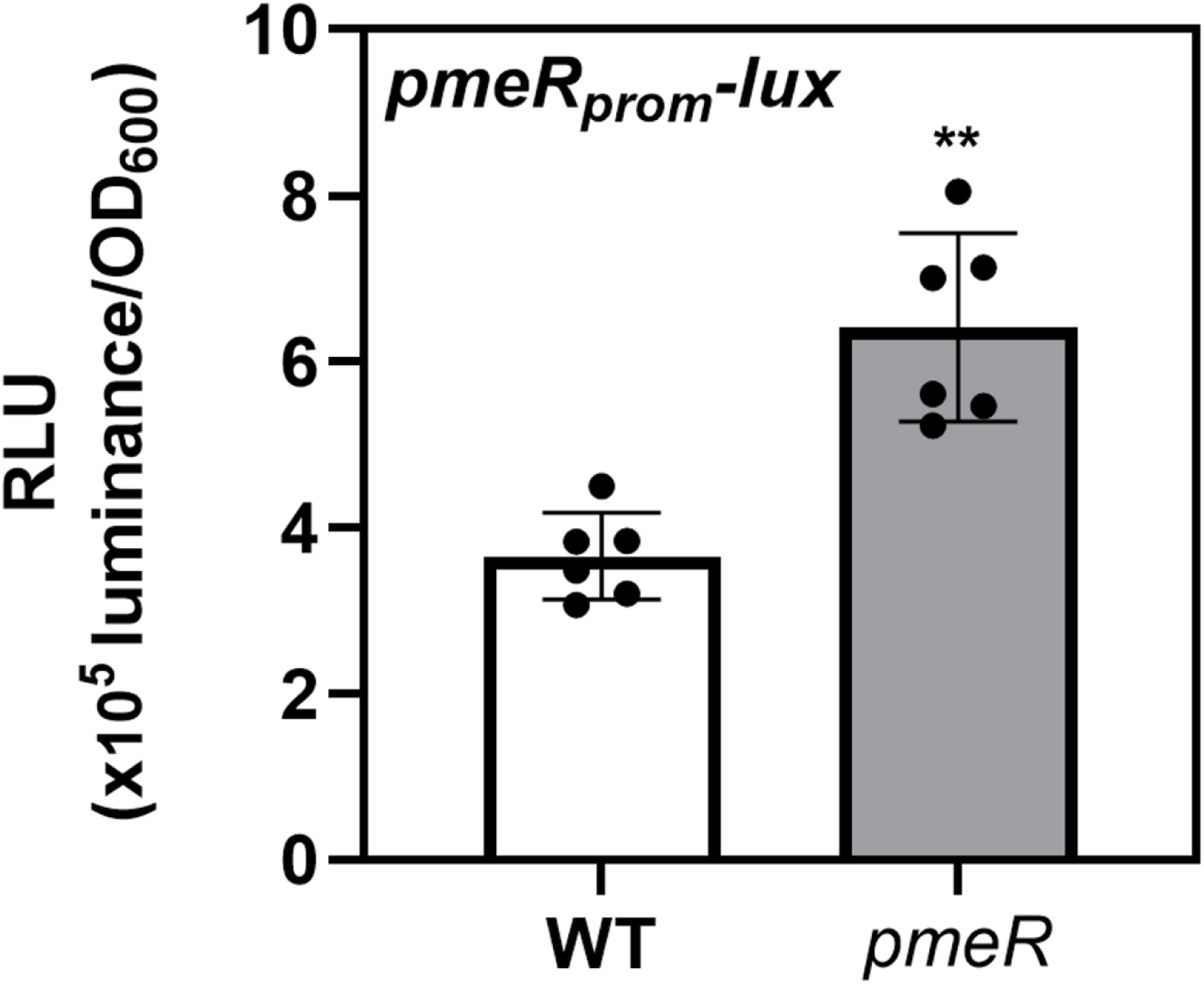
*pmeR* represses its own expression. Luciferase activity of the *pmeR_prom_-lux* reporter in wild-type *Pto*DC3000 (WT, white bar) and the *pmeR*::ΩKan mutant (*pmeR*, grey bar) in Hrp de-repressing medium (HDM). Data represents mean luminescence normalized to cell density (OD_600_), plotted as relative luminescence units (RLU) ± standard deviation (SD), and are compiled from two independent experiments (n = 6). Asterisks indicate significant differences between WT and *pmeR* mutant strains as determined by Student’s *t*-test (**: *p* < 0.01).

**Fig S2.**
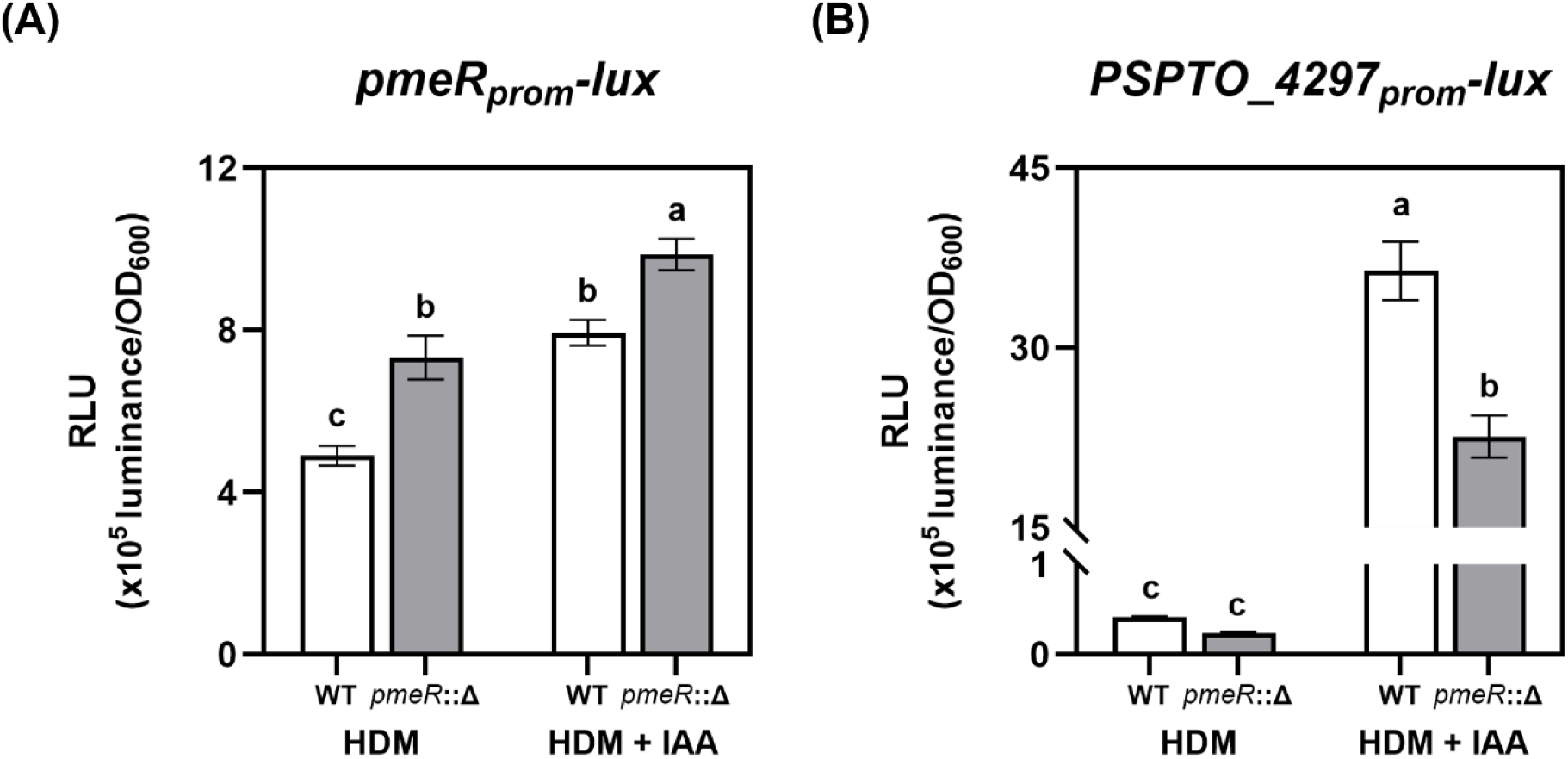
An independent deletion mutant of *pmeR* exhibits elevated expression of *pmeR* and reduced IAA induction of *PSPTO_4297*. Expression of (A) *pmeR_prom_-lux* and (B) *PSPTO_4297_prom_-lux* in wild-type *Pto*DC3000 (WT, white bar) and the *pmeR* deletion mutant (*pmeR*::Δ, grey bar) in response to IAA at 90 minutes post-treatment. Data are from a representative experiment (n = 6) and plotted as the mean of relative luminescence units (RLU) ± SD. Similar results were obtained in two additional independent experiments. Letters indicate significant differences between treatments as determined by ANOVA followed by Tukey’s HSD test (*p* < 0.05). HDM: Hrp de-repressing medium; HDM + IAA: HDM supplemented with 100 µM IAA.

**Fig S3.**
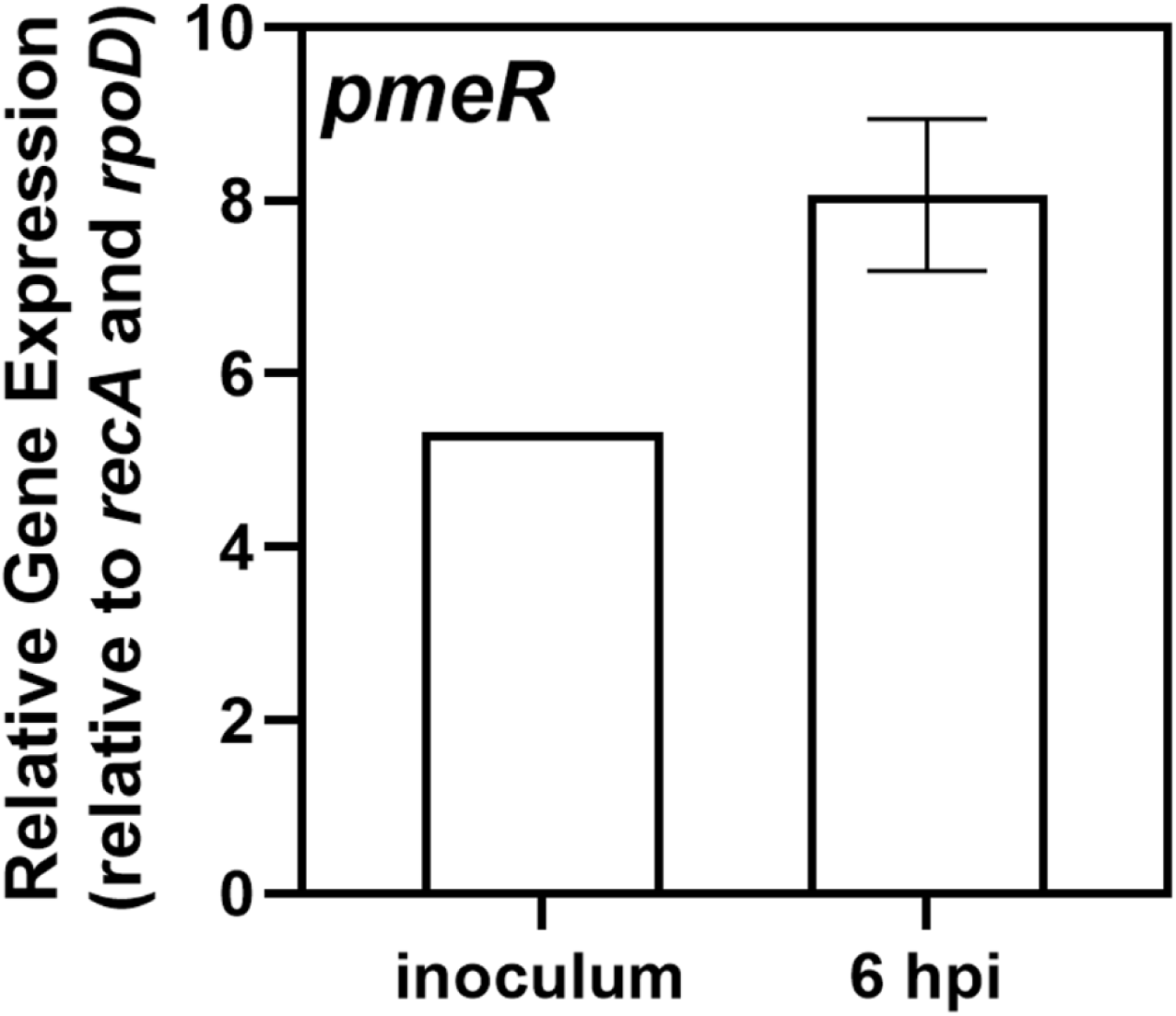
*pmeR* is expressed *in planta*. Expression of *pmeR* in *Pto*DC3000 growing in *A. thaliana* Col-0 plants. To quantify bacterial gene expression, infected leaves were harvested 6 hours after inoculation, and total RNA was isolated and used for qRT-PCR. Bacterial RNA from 1 mL of the initial *Pto*DC3000 cell resuspension was isolated and served as a control (inoculum, n = 1). The relative expression was calculated by normalizing *pmeR* expression to the expression of two reference genes, *rpoD* and *recA.* Data are from a representative experiment (n = 3) and plotted as the mean of relative gene expression ± SD. Similar results were obtained in a second independent experiment.

**Fig S4.**
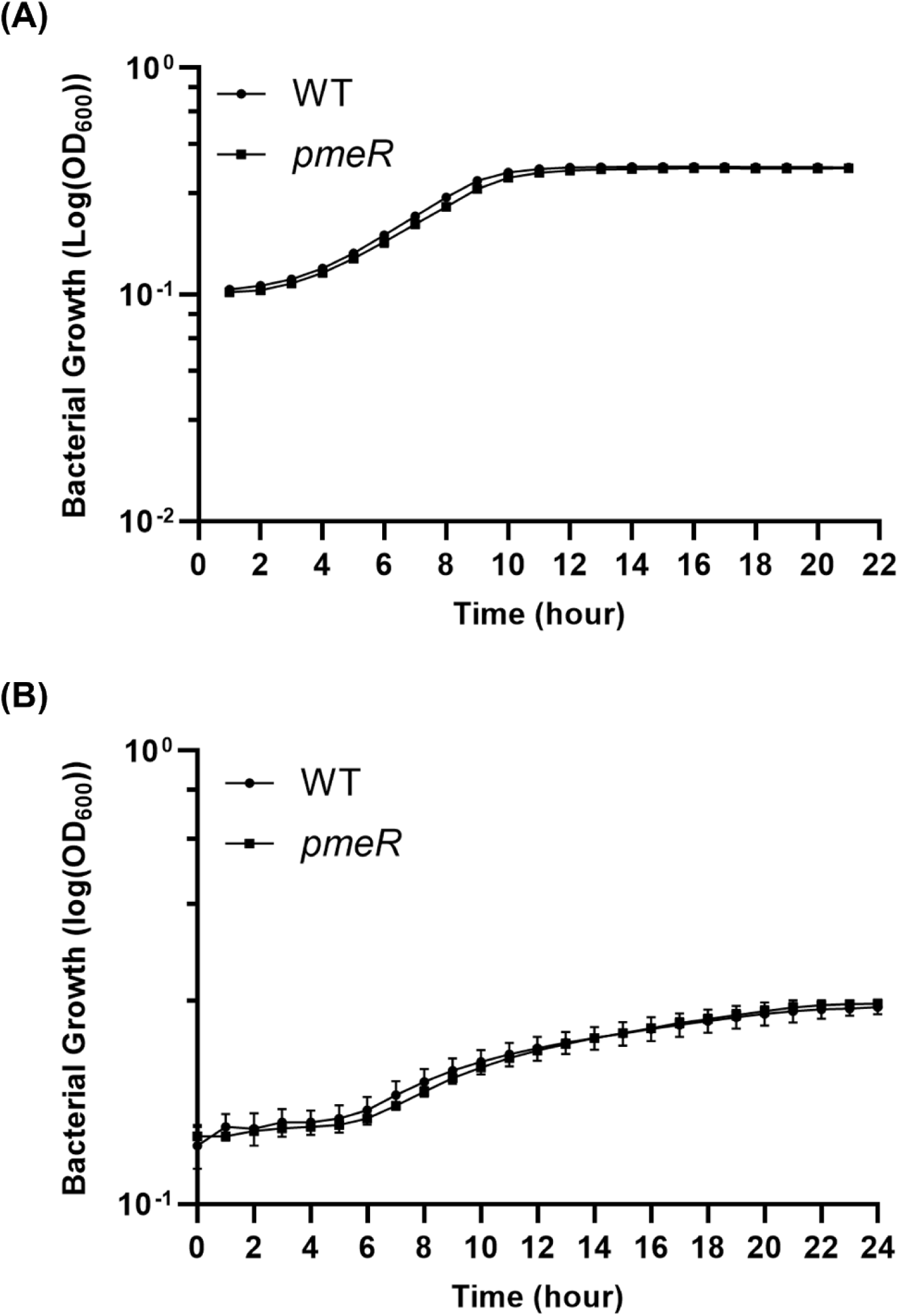
Growth of the *pmeR*::ΩKan mutant in culture. Bacterial growth of WT and *pmeR*::ΩKan (*pmeR*) in (A) rich (NYG pH 5.7) and (B) minimal media (HDM). Data are from one representative experiment and shown as mean ± SD (n = 2). No significant difference between strains was observed as determined by Student’s *t*-test. Similar results were obtained in an additional independent experiment.

**Fig S5.**
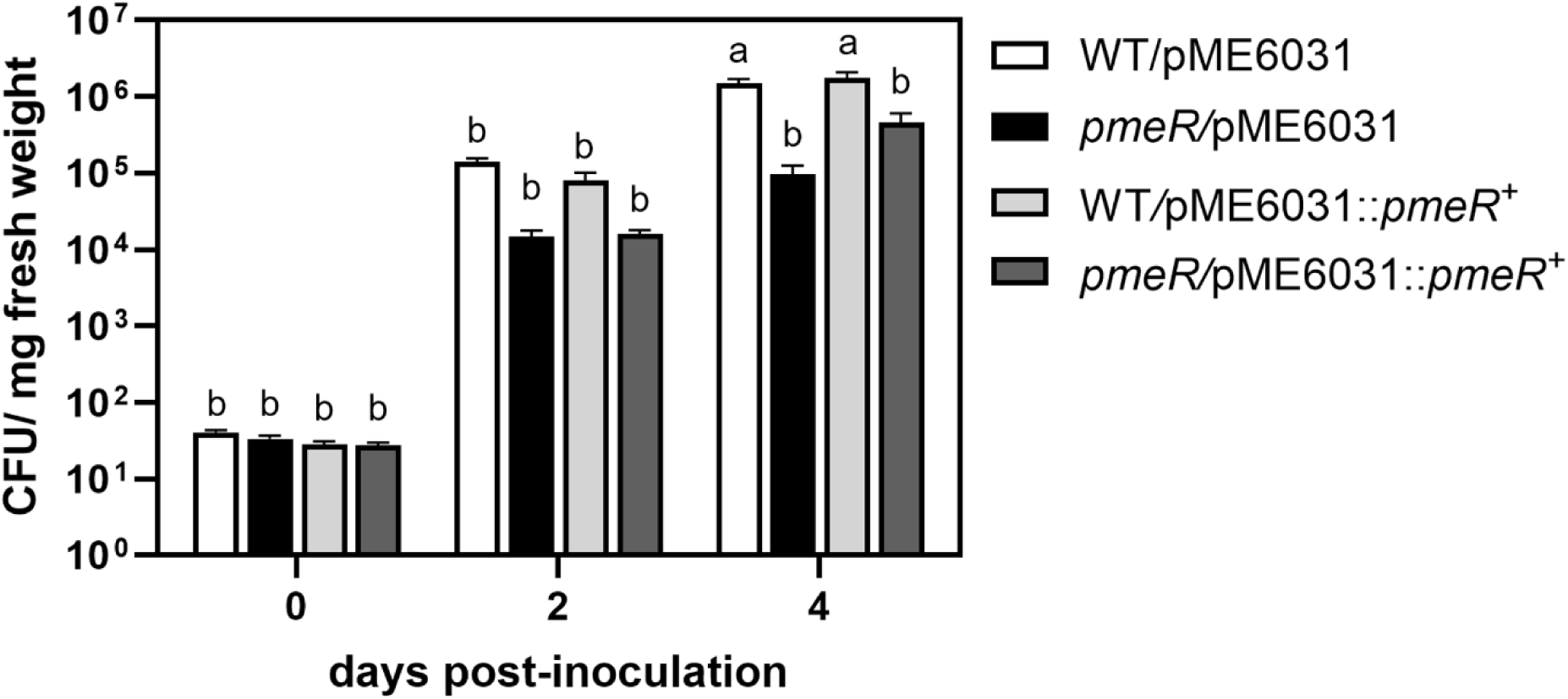
The wild-type *pmeR* gene partially complements the growth defect of *Pto*DC3000 in *A. thaliana*. Four-week-old wild-type *A. thaliana* Col-0 were infiltrated with ∼ 1 x 10^5^ CFU/mL of bacteria. Bacterial growth in infiltrated leaves was quantified at 0, 2, and 4 dpi. Data from two representative experiments are combined and shown as mean ± standard error (SE) (n = 8 for 0 dpi, n = 12 for 2 and 4 dpi). Letters indicate significant differences between treatments as determined by ANOVA followed by Tukey’s HSD test (*p* < 0.05). CFU: Colony forming units.

**Fig S6.**
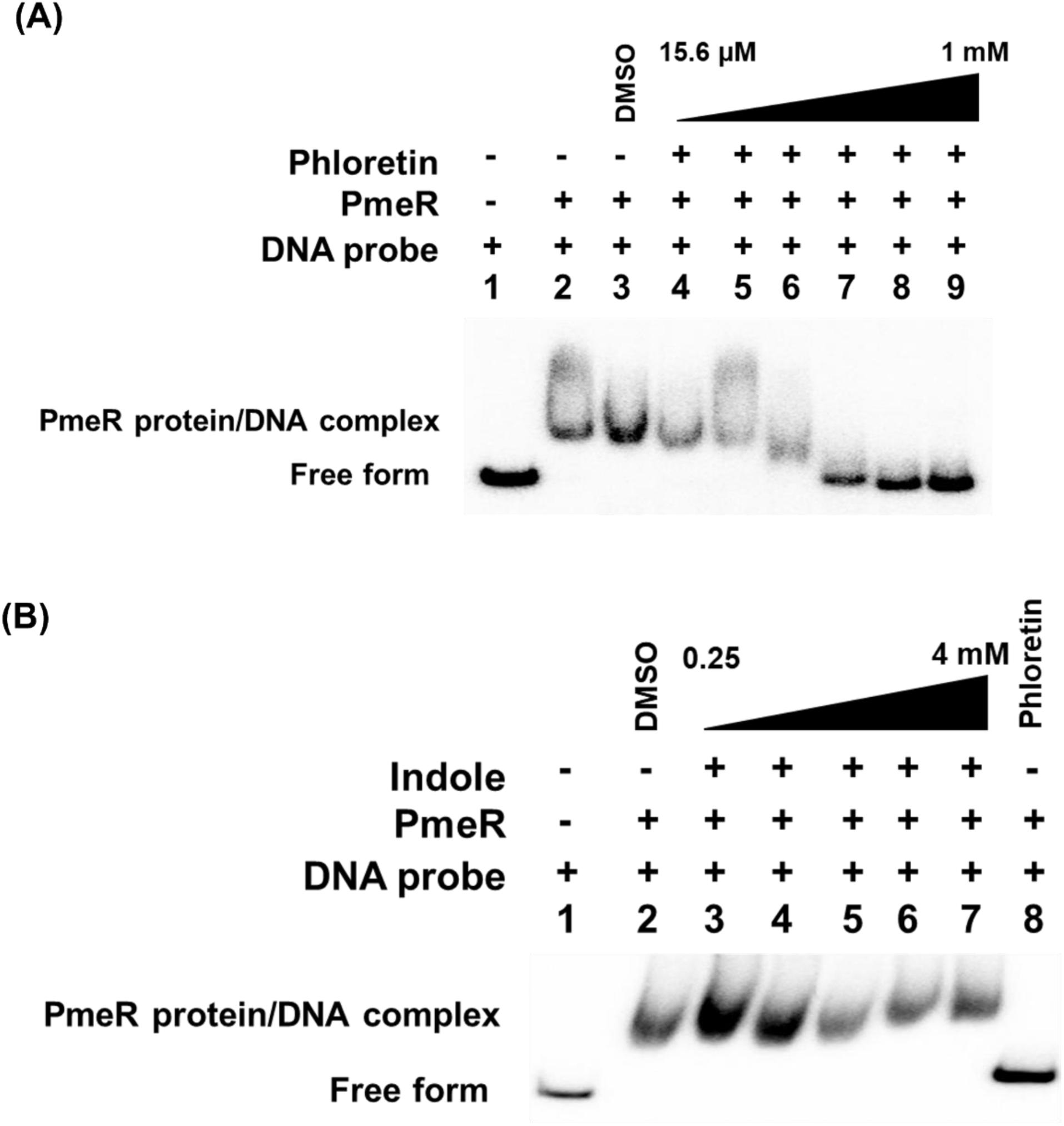
Phloretin disrupts the PmeR/DNA interaction. (A) The phosphor image of electrophoresis mobility shift assays (EMSA) confirms that phloretin disrupts the PmeR/DNA interaction. Increasing concentrations of phloretin (15.6 µM to 1 mM) were added to the co-incubation of 1.5 nM DNA probe and 6.25 nM purified PmeR protein. (B) The phosphor image of EMSA demonstrates that indole does not disrupt the PmeR/DNA interaction. Increasing concentrations of indole (0.25 mM to 4 mM) were added to the co-incubation of 1.5 nM DNA probe and 6.25 nM purified PmeR protein. For data shown in panels A and B, similar results were seen in two additional independent experiments.

**S1 Table. Bacterial strains and plasmids used in this study.**

**S2 Table. Primers used in this study.**

